# Computing tumor specificity of cancer antigen targets by k-mer indexing of healthy tissue transcriptomes

**DOI:** 10.64898/2026.06.29.734488

**Authors:** Johannes Hausmann, Franziska Lang, Özlem Muslu, Luis Kress, Jonathan Landry, Martin Suchan, Andrea Nubbemeyer, Ruprecht Kuner, David Weber, Barbara Schrörs, Marcel H. Schulz, Matthias M. Gaida, Ugur Sahin, Jonas Ibn-Salem

## Abstract

Individualized cancer immunotherapies rely on tumor-specific T-cell antigens, often predicted from somatic mutations as neoantigens. For tumors with low mutational burden, mRNA transcript variants, including gene fusions and novel splice junctions, can serve as important alternative targets. A main challenge in their identification from tumor RNA-seq is to confirm that their expression is tumor-restricted. Although large public collections of healthy-tissue RNA-seq exist, verifying tumor-specific expression requires computationally expensive re-analysis of these data for every novel candidate. To address this, we benchmarked nine k-mer indexing algorithms and developed k4neo, which leverages k-mer indexing of raw RNA-seq reads to compute the tumor specificity of any transcript variant. This mapping-free and transcript-class agnostic approach screens any candidate sequence against 18,960 samples across 51 healthy tissue types. We confirmed k4neo’s detection accuracy with qRT-PCR and showed that k4neo accurately classifies somatic and germline variants, gene fusions, and isoforms by tumor specificity. Applied to nine tumor cohorts, it nominated a median of 4-80 tumor-specific splice junctions per patient, including recurrent, long-read-confirmed novel antigen candidates. Together, k4neo enables efficient access to large-scale sequencing cohorts and accurately computes tumor specificity for any input transcript sequence, thereby expanding the repertoire of individual and shared cancer antigen targets.

## Introduction

Cancer immunotherapies rely on peptides presented by MHC molecules and recognized by T cells. These antigens fall into two classes: tumor-associated antigens (TAAs), which are self-proteins overexpressed in tumors, and tumor-specific antigens (TSAs), which are absent from the normal transcriptome and proteome. Because TAA-reactive T cells are subject to central and peripheral tolerance, TAA-directed responses are typically low-avidity and risk on-target/off-tumor autoimmunity, whereas TSAs can engage a high-avidity, tumor-restricted T-cell repertoire^1,2^

Most immunotherapies, including personalized cancer vaccines, target neoantigens arising from nonsynonymous somatic mutations, identified by comparing tumor and matched-normal DNA sequencing^3,4^. Tumor mutational burden (TMB), a proxy for neoantigen content, is correlated with clinical benefit of immune-checkpoint blockade across cancer types^5,6^ and personalized neoantigen vaccines induce CD4⁺ and CD8⁺ T-cell responses with clinical benefit in melanoma and other solid tumors^4,7–10^. However, many tumors have low TMB and yield too few mutation-derived targets^11–13^. Recent clinical trials show that neoantigen vaccination can succeed in low-TMB malignancies such as glioblastoma^8,14^ and pancreatic ductal adenocarcinoma^13^, but only when sufficient TSAs can be identified. Therefore, the discovery of tumor-specific non-canonical antigens has become a central bottleneck for individualized immunotherapies. The tumor-specificity requirement extends beyond vaccines: anti-body-, bispecific-, and CAR-/TCR-T-cell–based therapies likewise depend on antigens that are expressed in tumors but absent from critical healthy tissues, to avoid on-target/off-tumor toxicity^15^.

Non-canonical TSAs can be identified from non-canonical RNA transcripts. These include gene fusions^16,17^, alternative splice junctions^18–20^, intron retention^21^, exitrons^22^, cryptic ORFs^23–25^, and transcripts from transposable elements such as endogenous retroviruses^26–28^, all of which can be detected from tumor RNA-seq. For example, RNA-seq identified 500–24,000 novel splice junctions per melanoma sample, that were absent from GENCODE reference¹. However, confirming that such candidates are truly tumor-specific is difficult, because their expression varies across tissues and any single matched control (e.g., blood or tumor-adjacent non-neoplastic) is insufficient.

SNV-derived neoantigens can be confirmed as tumor-specific by matched-normal DNA sequencing, but no equivalent control exists for splicing- and gene fusion-derived peptides: their source transcripts are often expressed at low levels across healthy tissues, where even minimal off-tumor expression compromises immunogenicity and might lead to off-tumor toxicity^29,30^.

Across these antigen classes, tumor specificity is operationally defined as absence of the candidate sequence in panels of healthy reference samples, which most commonly consists of subsets of GTEx^31^, TCGA^32^ tumor-adjacent non-neoplastic samples, or the Human Protein Atlas, using filters that range from event-level statistics (e.g., PSI or z-scores)^19^ to binary absence calls by pre-build exclusion lists^21,27^.

While public resources for such comparisons are vast, e.g. GTEx^31^ alone contains >17,000 RNA-seq samples across 54 tissues, and the SRA exceeds 750,000 human samples^33^, exploiting them is challenging. Most filtering pipelines, including recent probabilistic frameworks that integrate gene- and protein-level expression^34^, operate on precomputed event tables (junctions, fusions, expression matrices) rather than raw reads, so a candidate detected in a new tumor cannot be re-evaluated without reprocessing thousands of normal samples. Sensitivity is further capped by choice of the aligner and computational pipeline as well as parameters such as reference genomes, transcript annotations and read-support thresholds. Events called “absent” may simply lie below the detection level, biasing calls toward false claims of tumor-specificity. Resources such as recount3^35^, which compiled gene, exon and junction counts from 316,443 samples, alleviate this for predefined event types but required ∼29,000 node-hours and 150 Tb of storage, and do not cover fusions, circular RNAs, or other variant classes. The alternative, to re-aligning all healthy reads against each candidate sequence^18,36^, is more sensitive but computationally prohibitive at the per-patient scale. A scalable solution should therefore evaluate any candidate transcript sequence against thousands of healthy RNA-seq datasets in a mapping-free, pipeline-independent manner, applicable to any class of non-canonical antigen.

Recent advances in k-mer indexing may facilitate the computationally efficient identification of TSAs.. Unlike transcriptome-indexing tools such as Kallisto^37^ or Salmon^38^, these algorithms index the raw reads of thousands of samples into compact data structures, by considering only short (19–31 bp) and unique subsequences (k-mer)^39–46^. While index construction is computationally expensive, it is precomputed once and subsequent queries return presence/absence efficiently^39^. However as the k-mer algorithms were mostly designed and evaluated for metagenomics studies^39^, it is unclear if they are sufficiently accurate and scalable to provide tumor-specificity assessment for antigen discovery.

Here, we benchmarked nine k-mer indexing algorithms and developed k4neo, a tool to compute the expression profile and tumor-specificity of any input transcript sequence across an index of 18,960 healthy RNA-seq samples spanning 51 tissues types. We show that k4neo accurately detects sample-specific splice junctions and predicts tumor-specificity for full-length tumor antigens, gene fusions, and somatic mutations. We applied it to nominate novel, highly tumor-specific splice junctions as antigen candidates across multiple tumor types, expanding the repertoire of safely targetable individual and shared targets for cancer immunotherapy.

## Material and Methods

### Benchmarking data for detection performance of splice junctions

To compare the accuracy of k-mer indexing methods to detect individual splice-junctions, we constructed a ground-truth dataset of canonical splice junctions. We used paired-end RNA-seq data from 148 samples of 48 different tissues (PRJNA764684)^36^. We identified expressed canonical splice junctions using a combination of read mapping against the hg38 reference genome using STAR^47^ and confirmation by a targeted mapping against the splice junction sequences with easyquant ^48^. More specifically, we used the splice junction annotations SJ.out.tab output of STAR (v2.7.10b) (with parameters `--runMode alignReads --alignIntronMax 500000 --alignMatesGapMax 500000 --outSAMtype None`), kept only canonical exon-exon junctions reported in GENCODE v40 annotations, and extracted the transcriptomic context sequences in a window of +/− 100 bp around each splice-junction using splice2neo^18^ (v0.6.2). We applied easyquant^18^ (v0.4.1) with options `-d 10` to re-quantify the number of supporting junction reads (mismatch free alignment in +/− 10 bp around the junction position) for all samples and all junction sequences. A sample-splice junction combination was classified as positive (expressed) if reported by STAR and detected by easyquant with at least two reads, as negative (not expressed) if the junction was not detected with STAR and easyquant. Other junction-sample pairs were not considered. The resulting 3,705 splice-junctions were grouped by the number of samples expressing them into sample specific (1 sample), rare (2-5 sample), medium (6-19 samples), and common (20-150 samples). To balance the dataset, we down-sampled the negative sample-junction pairs to the same number of positive sample-junction pairs per group, resulting in 76, 2875, 135, and 6732 positive and negative junction-sample pairs in the respective groups (Supplementary Fig. S1A). For every considered k-mer size in the benchmark, we shortened the transcript context sequences to 2*(k-1) bases around the splice-junction position.

### K-mer indexing methods

To determine the feasibility and accuracy of detecting novel RNA-derived transcript candidate sequences in k-mer indices, we selected and evaluated seven k-mer indexing methods. Five methods are based on approximate membership representations: Kmtricks (1.4.0)^40^, HowDeSBT (2.00.02)^41^, COBS (0.2.1)^42^, Raptor (3.0.1)^43^ and Kmindex (0.5.2)^44^. The remaining two are exact compact De Bruijn graph-based (cDBG) methods: MetaGraph (0.3.6)^45^ and REINDEER (1.0.2)^46^. For Raptor, we compared three k-mer indexing modes: the Interleaved Bloom Filter (IBF) data structure (Raptor_kk), indexing minimizers using IBF (Raptor_wk), or indexing minimizers with the Hierarchical Interleaved Bloom Filter (HIBF) data structure (Rap-tor_hibf_wk)^43^.

For all tools, we constructed indices for k-mers of size 19, 21, 25, and 31 bp. We removed k-mers that occur only once per sample, as these are likely of erroneous origin. If not noted otherwise, we used the tools with their default or recommended pre-processing strategies and parameters according to their documentation. For the Bloom filter (BF) parameter selection in Kmtricks, HowDeSBT and kmindex, we used ntcard^49^ to estimate the number of distinct k-mers in each cohort. For HowDeSBT and kmtricks, we set the size of the bloom filter to the estimated number of unique k-mers in the union of the experiments. For Reindeer and COBS, we used cDBG as input. For each sequencing sample, cDBGs were constructed with BCALM2^50^ while excluding unique k-mers. We choose cDBG as the input for COBS because this tool does not filter input FASTQ files, leading to the creation of extensively sized indices for sequencing collections. For MetaGraph, we constructed k-mer counters using KMC3^51^ and subsequently constructed the succinct de Bruijn graph representation. For Raptor, we set the window size for indexing with minimisers to 4 as suggested in the Raptor manual. For all tools, we build indices on a compute server with 96 CPU cores and 1.5 Tb of RAM. We parallelized the pre-processing steps of individual tools using GNU `parallel`^52^ where applicable to reflect typical real-world applications of the implementations.

### Calculating detection accuracy for k-mer indexing methods in benchmarking dataset

We queried the splice junction sequences from the ground-truth dataset in the indices using the default commands from the tools and considered a junction as detected per sample if a minimal fraction of k-mers from the query sequence (k-mer ratio) was found in the index for the sample. For all methods, k-mer sizes, and k-mer detection ratios (0.45, 0.5, 0.6, 0.7, 0.8, 0.9, and 1), we compared the detected sample-junction pairs with the ground truth data and calculated sensitivity (recall), precision (PPV) and F1-score as detection performance.

### Healthy tissue cohorts and SRA index building

To build a healthy tissue reference dataset, we initially collected studies with accessible RNA-seq data from as many samples and non-disease tissues as possible (Supplementary Table S4). We downloaded the raw RNA-seq data from SRA, GEO^53^ and ENCODE^54^. We applied fastp (v0.23.4)^55^ (with parameters `--stdout --unpaired1 --unpaired2 --length_required 31`) to remove low quality bases and adapter sequences. For paired-end sequencing data, we generated interleaved FASTQ files per sample, that include forward and reverse reads that pass QC filtering. We discarded the reads shorter than 31 bp (largest k-mer size used).

### Computational resource benchmarks

To assess the computational efficiency of k-mer indexing methods, we built indexes for subsets of varying sizes from the cohorts (Supplementary Table S1, S2) on a single cluster node using 16 CPU cores for each job and a k-mer size of 21 bases^41,56^. We recorded the run time, CPU usage, peak memory usage and file size of the final index using GNU time and psrecord (Supplementary Table S1). In contrast to previous work, we excluded only unique k-mers, without applying the low-frequency k-mer threshold used in previous publications^41,46,57^. We built two independent sets of query sequences by randomly selecting 10,000 canonical splice junctions from the GENCODE v40 reference transcriptome twice and annotating them with +/− 50 bp of flanking transcript sequence. We assessed the required query time and peak memory usage for querying the SRA k-mer index from each tool. We successively queried the two sequence sets to reflect cold and warm caching scenarios.

### k4neo

We developed and implemented the tool k4neo (https://github.com/TRON-Bioinformatics/k4neo) to annotate any input transcript sequence, e.g. novel splice junctions, with expression profiles in healthy and cancer tissue samples. k4neo is implemented as a python package and relies on the k-mer indexing methods Raptor^43^ or kmindex^58^. We provide the pre-built SRA k-mer index (k=21, as determined in our benchmark) with k4neo for 2,256 manually curated RNA-seq samples of 49 healthy tissues from public resources (Supplementary Table S4). Curated metadata per sample, such as tissue, developmental state, disease and optionally sex and age are stored in a document-oriented TinyDB database. In search mode, k4neo accepts transcript sequences as input and optionally a position of interest (e.g. splice junction or gene fusion breakpoint). These query sequences are then decomposed into overlapping k-mers, which are searched with a given k-mer ratio parameter in the pre-built indexes by Raptor or kmindex. The matches in k-mer indices are parsed as pairs of input sequence and individual RNA-seq sample and annotated with tissue, disease and developmental state information. Finally, for each input sequence, k4neo computes expression profiles as sample rate (fraction of samples for which the sequence was detected) per annotated metadata features (tissue, developmental state and disease). In addition, k4neo can be used in build mode to construct k-mer indices using Raptor and kmindex for additional samples from RNA-seq raw data.

### Indexing healthy tissue samples from GTEx and tumor samples from TCGA

To index healthy tissue samples and pan-cancer tumor samples for k4neo, we selected healthy tissue samples from GTEx v9 and samples from TCGA^31,32^. From GTEx samples we excluded cell lines and cultured fibroblasts totaling 16,704 samples for the GTEx index (Supplementary Table S4). For the TCGA we indexed tumor (n=10,328) and adjacent normal samples (n=737) (Supplementary Table S4). Indexed samples and associated metadata can be found under https://github.com/TRON-Bioinformatics/kmer_index_data. We developed a Workflow Description Language (WDL) script to generate k-mer indices from aligned RNA-seq BAM files using Raptor v3.0.1 in TERRA bioinformatics cloud (app.terra.bio). We constructed Raptor indices using the HIBF layout and parameters (=21, =25) as determined in our benchmark. As GTEx and TCGA raw data is only available under restricted access we do not provide the pre-build index, however, we provide WDL and Snakemake workflows for users to construct easily custom k-mer indices that are directly usable with k4neo.

### Claudin 18 isoform analysis

We selected the isoform 1 and 2 of the gene Claudin 18 (CLDN18), which differ in their first exons and are differentially expressed in samples of the lung and stomach^59^. We represented each transcript variant by +/− 20 bp sequence around the respective first splice junction (Supplementary Table S5) and queried it against the GTEx/SRA tissue index using k4neo. For comparison, we downloaded raw STAR splice-junction counts from the Genotype-Tissue Expression (GTEx) portal (`bulk-gex_v8_rnaseq_GTEx_Analysis_2017-06 05_v8_STARv2.5.3a_junctions.gct`)

### Cancer testis antigen analysis

To assess the ability of k4neo to detect the disease-specific expression patterns of full-length shared antigen candidates, we queried the representative transcripts (MANE or ENSEMBL canonical) of 18 cancer testis antigens (Supplementary Table S6) against healthy tissue and cancer k-mer indices. Besides the healthy tissue index we analyzed melanoma patients with RNA-seq samples from Hugo et al.^60^ (n=27, SRA SRP070710) and Riaz et al.^61^ (n=46, SRA SRP094781) with k4neo as described above. For comparisons, we calculated gene expression in transcript per million (TPM) with kallisto (v0.46.0)^37^ for the melanoma samples.

### qRT-PCR confirmation

To assess the ability of k4neo to detect splice junction sequences in RNA-seq using parameters optimized on the in-silico ground-truth dataset, we generated an orthogonal benchmark based on qRT-PCR validation. We used RNA-seq data from the MCF7 and SKBR3 cell lines, previously generated at our institute^16^, for this analysis. We selected 50 expressed canonical splice junctions (25 per cell line) as positives following the same criteria used for the in-silico benchmark. The negative set comprised 25 junctions from non-expressed genes (kallisto TPM = 0) and 25 randomly generated exon-exon fusion junctions without any read support. Primer design and qRT-PCR experiments were performed as described previously^18^. Amplicons were evaluated using capillary gel electrophoresis (QIAGEN’s QIAxcel Advanced), with ambiguous results resolved by Sanger sequencing. Based on qRT-PCR and Sanger confirmation, junction–cell line pairs were classified as positive or negative, resulting in 48 positives and 68 negatives (Supplementary Table S9). We indexed the MCF7 and SKBR3 cell line RNA-seq data (SRA PRJNA543964) with k4neo and queried these sequences against the k-mer index, to assess detection performance in terms of sensitivity and specificity.

### Germline and somatic mutations in melanoma

To test whether k4neo can predict the tumor specificity of point mutations, we assessed whether the sample rate across healthy tissue samples distinguishes somatic single-nucleotide variants (SNVs) from germline single-nucleotide polymorphisms (SNPs). We analyzed whole-exome sequencing (WES) and RNA-seq data of n=73 melanoma samples^60,61^ as described previously^18^. For somatic variants WES of tumor and normal was aligned against hg19 reference genome using bwa (v0.7.17)^62^ and subsequently somatic variants were called using Mutect2^63^ in the TronFlow workflow collection pipelines^64,65^. Germline single nucleotide polymorphisms (SNPs) were detected by applying the tronflow-haplotype-caller pipeline (v1.0.1)^66^ which is based on GATK (v4.2.6.1) on aligned normal sequencing data, restricting detection to exonic target regions and applying filtering with variant quality score recalibration using dbSNP^67^, the 1000 Genomes Project^68^ and HapMap^69^ sets of high quality known variant sites. Germline variants were then annotated with gnomAD^70^ population allele-frequency and filtered for a minor variant allele frequency (MVAF) >= 0.05. Next, germline and somatic variant calls were inserted into coding transcript sequences from GENCODE (v45lift37) annotation and shortened to +/−20 bp around the mutation. If multiple mutated transcripts generated the same context sequence, we collapsed them into one search query (Supplementary Table S10). Subsequently, context sequences were annotated with expression profiles in the GTEx/SRA index using k4neo.

### Analysis of gene fusion transcripts associated with structural variants

To compare the tumor-specificity of gene fusions associated with structural variants (SVs) versus RNA-only detected events, we reanalyzed CCLE data from 327 cell lines^71^. We used EasyFuse v2.0.4 to predict gene fusions events from RNA-seq data (model prediction score >= 0.5). Somatic SVs identified from whole genome sequencing (WGS) via SvABA were obtained from DepMap Portal, were lifted from GRCh37 to GRCh38 with rtracklayer^72^ and annotated with overlapping genes using Ensembl v89 reference annotation. RNA-seq fusion transcripts were matched to SVs based on gene pairs and breakpoint proximity (≤100 kb). Gene fusions for which both gene partners overlapped at least one SV were classified as “with SV”, those with only one gene partner affected (“partial SV”) were excluded, and gene fusions with no overlapping SVs were classified as “without SV” (Supplementary Table S11). The fusion transcript sequences (context sequence around breakpoint) were searched with k4neo against the GTEx/SRA index.

### Re-analysis of mutation-induced splice junctions in melanoma

We used splice junction data from RNA-seq of n=73 melanoma samples^60,61^, reported in previous work^48^, where we predicted tumor-specific splicing in these samples with the tool splice2neo by associating the effect of somatic SNVs in cis-regulatory regions (e.g. gain or loss of splice motifs) with splice junctions identified in tumor RNA. In short, the effect of a mutation on splicing was predicted using SpliceAI (v1.3.1)^73^ and MMSplice (v2.1.1)^74^ using default parameters^60,61^. Mutation-derived splice junctions were filtered with absolute prediction scores 0.35 returned by MMSplice and SpliceAI. Expressed splice junctions were predicted using Spladder (v3.0.2) ^75^ and Leafcutter (v0.2.9)^76^ based on the spliced alignment of STAR (v2.7.0a) against the hg19 reference genome. The R-package splice2neo was then used to combine the RNA-derived and mutation-associated splice junctions and subsequently annotated them with the corresponding transcript sequence. This analysis resulted in 59 mutation-associated junctions. Here, we additionally analyzed the purely RNA-derived junctions that were not associated with a mutation. We considered non-canonical junctions that were absent from an exclusion list based on GENCODE reference annotation and healthy samples from GTEx^48^, resulting in 33,552 RNA-derived junctions. We queried all junction sequences with k4neo in the GTEx/SRA healthy tissue index and compared the number of junctions per tumor sample and junction class that were detected with different tumor-specificity filter criteria: in zero, at least one percent or more healthy tissue samples.

### Splice junction detection from RNA-seq data

We developed and implemented a computational pipeline, called NeoRasp, to identify tumor-specific splice junctions from RNA-seq data from individual tumor samples (Supplementary Figure S5). NeoRasp incorporates state-of-the-art tools for splicing prediction tools and is implemented as reproducible Snakemake (v9.13) ^77^ workflow with software dependencies managed with Open Container Initiative (OCI) containers and Apptainer. Given paired-end FASTQ files as input, reads are aligned against the reference genome (here hg38) using STAR (v2.7.11)^47^ using ENCODE3 parameters (see STAR manual). Splice junctions from the high confidence “SJ.out.tab” are filtered based on a user-defined detection cut-off (here >= 5 uniquely mapped reads). Canonical splice junctions from the reference gene annotation (here GENCODE v46) are removed from the detected candidates. Next, the R-package FRASER (v2.0)^78^ is used to calculate the splice junction usage (Intron Jaccard Score) of each candidate junction using the genomic alignment. Splice junctions are annotated with counts per million (CPM) normalized read count values and reference gene and transcript expression values (TPM) estimated with Salmon (v1.10)^38^ on the STAR alignment in transcript coordinates (Aligned.toTranscriptome.out.bam). Subsequently, splice2neo (v0.6.13)^18^ is used to annotate junction candidates with corresponding transcript and peptide sequence. The read support of candidate splice junctions is then re-quantified in a targeted manner with easyquant (v0.6.0)^18^ using the transcript context sequences around the junction with stringent bowtie2 (v2.5.3)^79^ alignment parameters (`--no-mixed --dpad 0 --gbar 99999999 --mp 1,1 --np 1 --score-min L,0,-0.01`). In the final step, all annotations and scores of the junctions are combined into a unified table. For splice junctions leading to change in the peptide sequence, a separate output file is generated that is compatible with NeoFox^80^ to annotate MHC-I and MHC-II binding epitopes and further (neo)antigen features.

### Individual and shared tumor antigen candidates from RNA-seq derived splice junctions

To identify individual and shared tumor antigen candidates from splice junctions, we predicted splice junctions detected in tumor RNA-seq using the previously described NeoRasp pipeline. We collected studies with accessible RNA-seq data from tumor-entities with low mutational burden and downloaded the raw FASTQ data from SRA (Supplementary Table S7). We gathered the results of individual samples into a unified table and annotated the junctions for tumor specificity using k4neo.We defined splice junctions with the following per sample criteria as reliable events: (i) 5 uniquely mapped reads; (ii) Junction read support CPM >= 0.1; (iii) Splice usage (Intron Jaccard) >= 5 %. We defined splice junction as shared events in a tumor cohort when it was detected in 5% of the samples of a tumor cohort but at least in 5 samples and reliably expressed in at least 1% of the cohort but at least two samples. As control we applied the same methods to a set of independent healthy tissue RNA-seq cohorts (Blood n=99^81^, Brain n=83^82^, Heart n=40^83^, and Liver n=26^84^) which are not included in the GTEx/SRA k-mer index. Next, we filtered out splice junctions that did not alter the peptide sequence of any transcript and splicing events that involved deep intronic splice-sites (> 50bp away from next canonical donor/acceptor) for which the reference-based peptide annotation was not unambiguously possible.

### HLA-typing and MHC-binding prediction

HLA-types of individual tumor samples were determined with seq2HLA (v2.3)^85^ based on RNA-seq data using default parameters. HLA-types of individual samples were collected into a unified table following the NeoFox^80^ patient-data format (see NeoFox manual). Next, NeoFox (v1.2.2)^80^ was used to annotate splicing derived peptide sequences with MHC binding prediction using default parameters. MHC binding predictions were classified into strong and weak binders using the following cut-offs^86^.

- MHC-I strong binder: NetMHCpan_bestRank_rank <= 0.5
- MHC-I weak binder: NetMHCpan_bestRank_rank <= 2
- MHC-II strong binder: NetMHCIIpan_bestRank_rank <= 1
- MHC-II weak binder: NetMHCIIpan_bestRank_rank <= 5

### Long-read confirmation of splice junction candidates

To confirm the existence of the short read predicted splice junctions we used matched and unmatched Oxford Nanopore (ONT) sequencing data for 19 UVM samples^87^, five SCLC cell lines (H526, H211, SHP77, H146, H69)^88^ and one AML sample^89^. We aligned raw reads with minimap2^90^ against HG38 using GENCODEv46^91^ as reference (with options: -`ax splice --secondary=no --junc-bed gencode_46.bed)`. We extracted splice junctions and annotated from generated alignment using regtools^92^ (with options: `regtools junctions extract -a 12 -s XS -t ts <in.bam> | regtools junctions annotate -o <out.junc> - <hg38_primary_assembly.fa> <gencode46.gtf>.)` We searched junctions within generated BED12 files and junctions supported by two or more long-reads were considered confirmed.

## Results

### k-mer indexing can accurately detect splice junctions but methods differ in computational scalability

The computational efficiency of k-mer indexing methods derives from strong compression of raw sequencing data, achieved by storing only nonredundant k-mer sequences per sample^39^. This compression raises the question of whether such methods retain sufficient sensitivity to detect query sequences within the indexed data. We therefore evaluated whether short query sequences, such as splice junctions, can be reliably detected, even when they are expressed in only a small subset of samples. We constructed a benchmark dataset from RNA-seq data of 148 samples across 48 healthy tissues^16^, comprising 3,705 unique canonical splice junctions. Using stringent alignment-based re-quantification, we defined 271,275 expressed (positive) and 274,203 non-expressed (negative) sample-junction pairs (Fig. 1A). Junctions were grouped into four classes based on the number of expressing samples (common, medium, rare, and sample-specific), which correlated with junction read support (Supplementary Fig. S1A,B).

**Figure 1.**
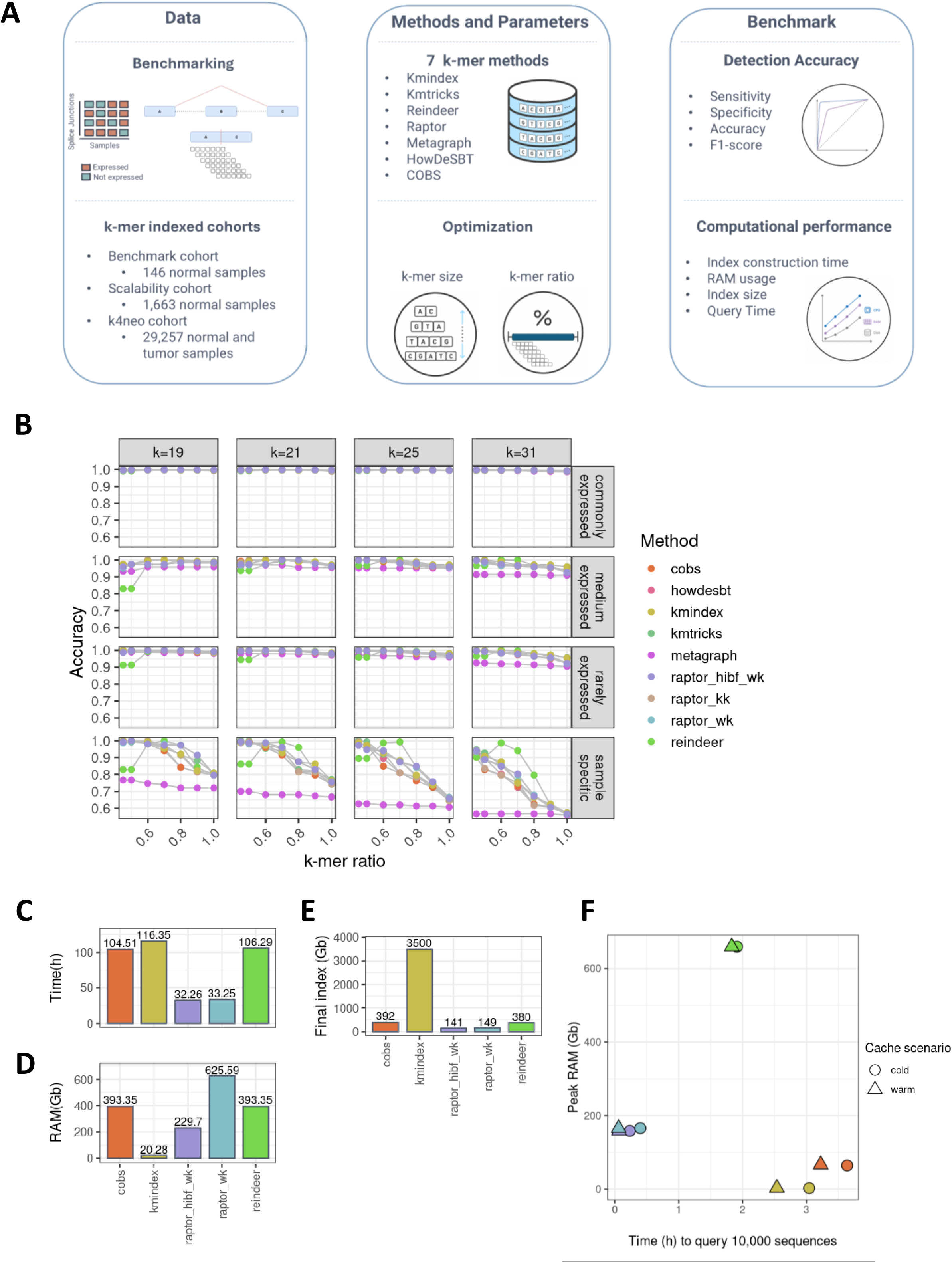
Benchmarking methods to index raw RNA-seq data for their accuracy in detecting splice junctions and their computational efficiency. **(A)** Overview of benchmark and method optimization. The benchmark dataset consists of 3,705 canonical exon-exon junctions. For each of 148 RNA-seq samples, each junction was labeled positive if at least one read was identified that support the splice junction by read mapping with STAR and easyquant and negative, if both methods did not find any supporting read. Splice junction sequences were split into different expression classes according to number of detecting samples. K-mer indexing methods were compared for their detection accuracy and computational resources in building indexes and searching in them. **(B)** Detection accuracy by method for multiple expression classes of splice junctions (rows) and different parameters for k-mer size (columns) and k-mer ratio thresholds (x-axis). **(C)** Wallclock time in hours by method to construct k-mer index from raw RNA-seq data. **(D)** Peak memory usage in Gb during index construction. **(E)** Final index size in Gb on disk. **(F)** Performance to query 10.000 canonical splice junctions in the index as peak RAM by query time for the methods Raptor, Reindeer, Kmindex and COBS for cold and warm (consecutive executions) cashing scenarios.

Using this benchmark data, we systematically evaluated seven k-mer indexing methods (Kmtricks ^40^, HowDeSBT ^41^, COBS ^42^, Raptor ^43^, Kmindex ^44^, MetaGraph ^45^, and REINDEER ^46^) across multiple parameters for k (k-mer size) and required fraction of detected k-mers per query sequence (k-mer ratio). For commonly expressed junctions, all methods achieved nearly 100% accuracy across all parameters (Fig. 1B). For medium and rarely expressed junctions, the detection accuracy decreased with increasing k-mer size and k-mer ratios down to 90%. Nevertheless, for specific parameters, all methods achieved accuracy between 95% and 100% on these junctions (Fig. 1B). Next, we focused on the more challenging sample-specific junctions that are expressed in only a single sample. All methods except for Reindeer reached specificity of more than 97%, and except Reindeer had no false-positive predictions across all parameter settings (Supplementary Fig. S1C). The sensitivity decreased with increasing k-mer sizes and ratios, most prominently for MetaGraph, which showed consistently lower performance across all tested parameters. All remaining methods achieved 95–100% accuracy for sample-specific junction detection when the optimal k-mer size and ratio were used (Fig. 1B).

After establishing detection accuracy, we benchmarked a subset of methods for scalability and computational efficiency using five batches of RNA-seq datasets, ranging from 122 to 1,663 samples (0.7–13.51 Tbp; Supplementary Table S2). When building indexes of increasing cohort sizes, we observed that HowDeSBT, kmindex, MetaGraph, and Raptor canonical k-mer mode exhibited exponential growth in either runtime or memory consumption (Fig. 1C-E; Supplementary Fig. S1E,F). Kmindex produced 10–25-fold larger indices than other methods, but required the least memory (Fig. 1D,E; Supplementary Fig. S1F,G). Raptor minimiser-based modes indexed the largest dataset in 32.26 h while producing the smallest indices (141 Gb). The HIBF variant required substantially less memory than the minimiser mode using canonical k-mers (229.7 Gb vs. 625 Gb). We evaluated the search time while querying two random sets of 10,000 splice junction sequences against the largest index (Fig. 1F). Raptor minimiser modes were fastest in both cold- and warm-cache scenarios, with query times of 14–24 min and around 4 min, respectively (Fig. 1F).

In summary, all evaluated methods achieved high sensitivity and specificity for detecting broadly expressed splice junctions, and, except for MetaGraph, accurately identified sample-specific events with their optimal k-mer size and rate parameters. The HIBF implementation in Raptor with k-mer size 21 and k-mer rate 70% provided the best balance of accuracy, computational scalability, and query performance. We therefore decided to use it in the implementation of k4neo to enable large-scale indexing for tumor-specificity prediction.

### k4neo computes expression profiles across large cohorts of healthy tissue samples and tumor types

To compute expression profiles and assess the tumor specificity of any input transcript sequence, we developed k4neo, a Python package that screens a query sequence against pre-built k-mer indices of raw RNA-seq reads (Fig. 2A). Unlike map-ping- or annotation-based approaches, k4neo is alignment-free and agnostic to the class of transcript variant, allowing, for example, junctions, gene fusions, isoforms, or point mutations to be evaluated against thousands of unmatched samples without a precomputed, event-specific reference table or exclusion list. For a given query, k4neo decomposes the input sequence into overlapping k-mers, searches them against the indexed samples, and integrates each match with the corresponding sample metadata. From these matches, k4neo computes expression profiles as the sample rate (the fraction of samples in which the sequence is detected) across healthy tissue groups and tumor types. In addition to querying pre-built indices, k4neo supports a build mode to construct custom indices directly from RNA-seq data.

**Figure 2.**
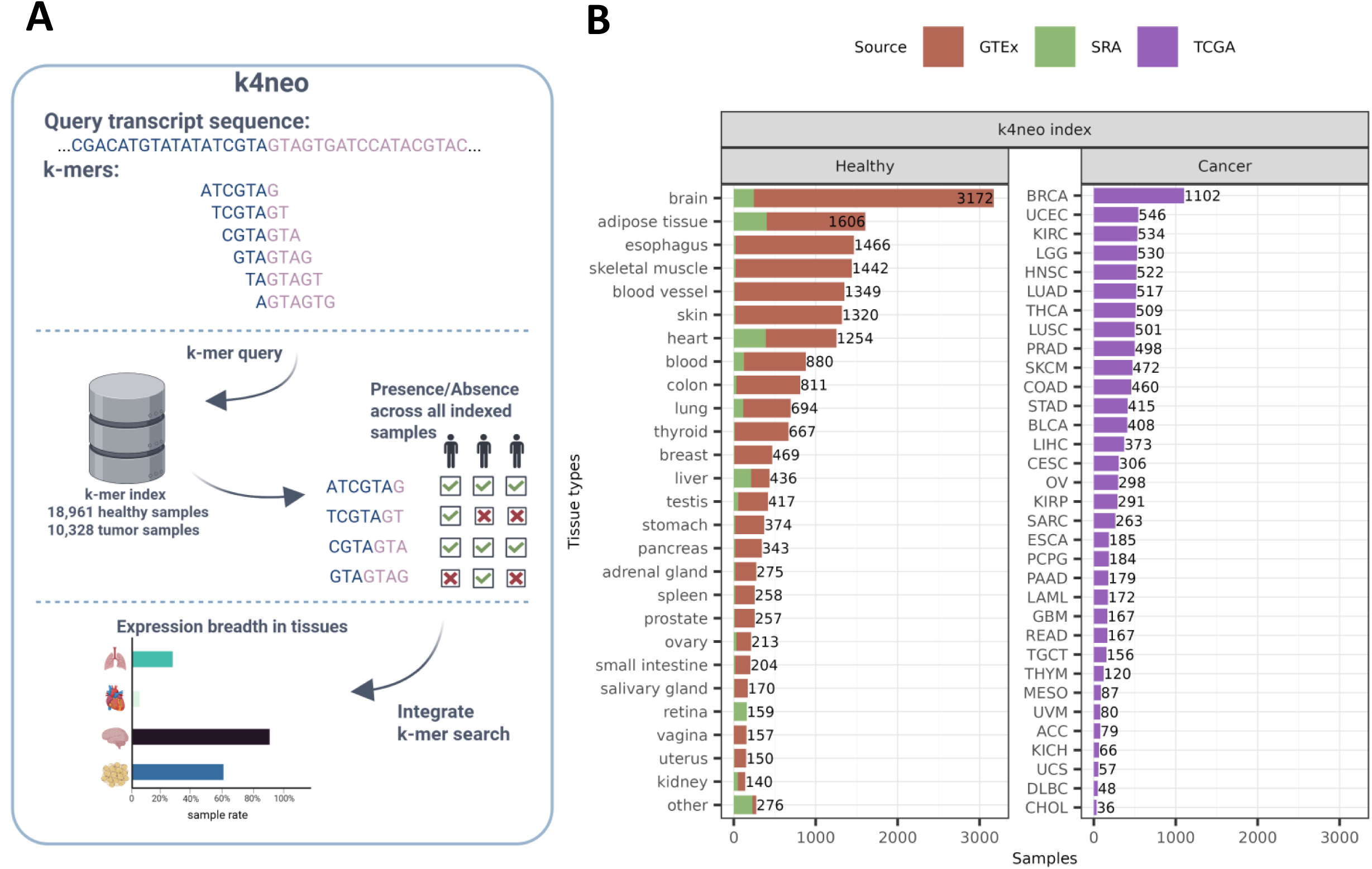
For any input sequence k4neo computes expression profiles across a diverse collection of healthy tissues and tumor types. **(A)** Overview of functionality implemented in k4neo python package. k4neo uses pre-built k-mer indices of RNA-seq samples. The input query transcript sequences are decomposed into overlapping k-mers, which are searched in the pre-built indexes. The matches of input sequence and samples are parsed and annotated with sample metadata. k4neo computes expression profiles as sample rate (faction of samples for which the sequence was detected) per annotated metadata features (tissue, developmental state, disease, and cancer type). **(B)** Number of RNA-seq samples in the k4neo indexed per healthy tissue (left) and tumor types from TCGA (right).

To provide broad reference coverage for k4neo, we aimed to maximize the number of human tissue types and samples with available raw RNA-seq data (Fig. 2B). We incorporated 16,704 samples from GTEx v9^93^ (GTEx index), and extended this resource with 21 additional manually curated studies, comprising 2,256 healthy tissue samples (SRA index). The SRA index added 23 tissue types absent from GTEx v9 and increased coverage of tissue types underrepresented in GTEx (Fig. 2B, Supplementary Fig. S2). Tissue names in the SRA index were harmonized to GTEx conventions and refined with sub-tissue types where available, yielding a total of 51 tissue types, 28 of which matched GTEx definitions. In total, this comprised 18,960 healthy RNA-seq samples, exceeding the sample sizes used for previously reported exclusion lists based on GTEx v3^73^ or GTEx v6^94^. To extend k4neo to tumor samples, we additionally indexed 10,328 tumor and 737 tumor-adjacent non-neoplastic samples from TCGA, spanning 31 tumor types (TCGA index; Fig. 2B). This allows any candidate sequence to be queried not only for healthy-tissue expression but also for its recurrence and expression across tumor types.

To validate k4neo’s detection accuracy experimentally, beyond the *in silico* benchmark of individual indexing methods, we performed a targeted qRT-PCR assay with Sanger confirmation for selected splice junctions in the breast cancer cell lines MCF7 and SKBR3^16^. This yielded 48 confirmed-positive and 68 confirmed-negative junction sequences. Using k4neo, we build an k-mer index from RNA-seq of these cell lines and queried the junction sequences. k4neo detected 45 of 48 confirmed-positive junctions (93.8% sensitivity), excluded 67 of 68 negative junctions (98.5% specificity), and achieved 96.6% overall accuracy (Fig. 3A).

**Figure 3.**
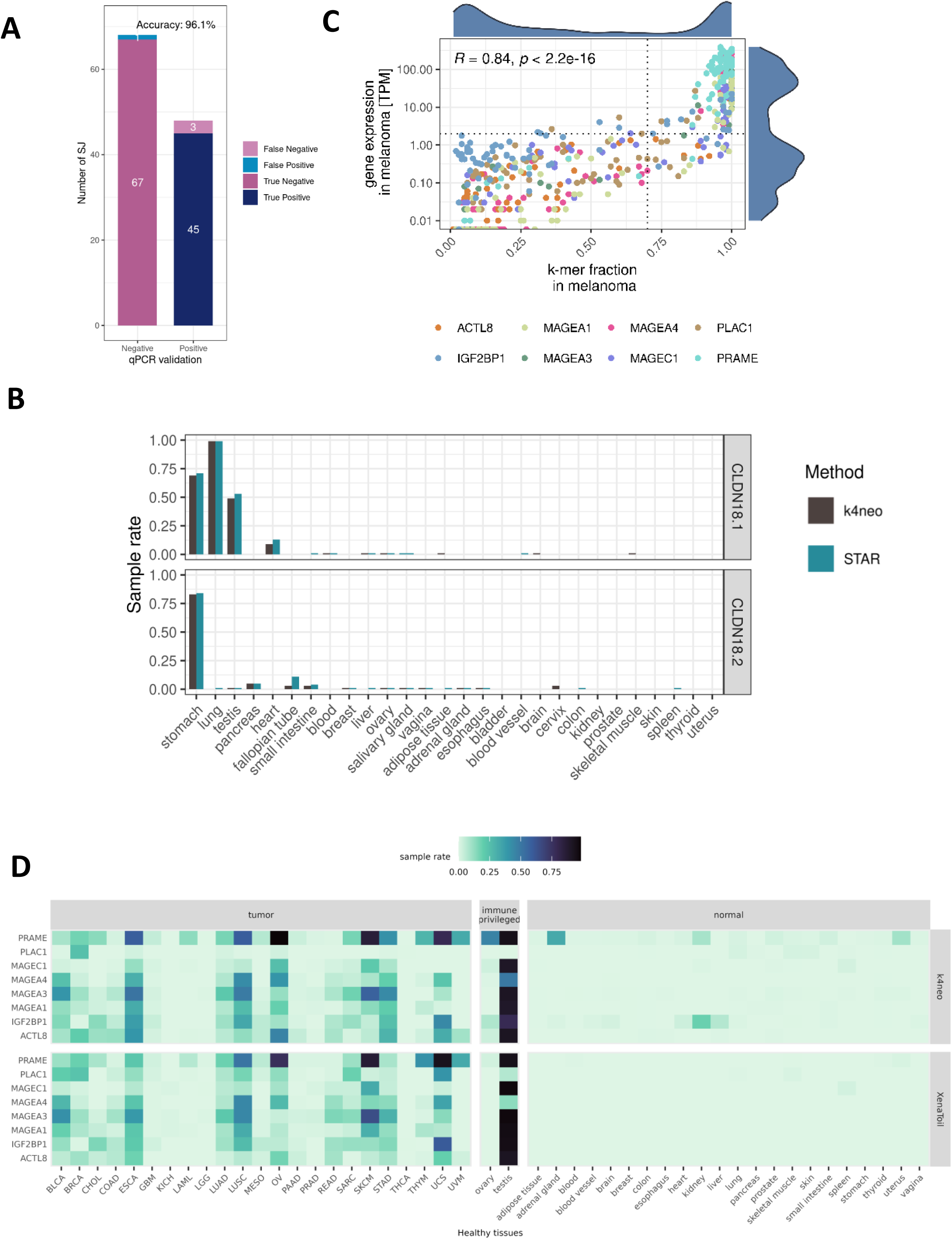
k4neo captures tissue-specific expression patterns of isoforms, full-length genes, and known cancer testis antigens. **(A)** k4neo detection performance on an orthogonal generated qRT-PCR splice junction (SJ) benchmark dataset. **(B)** Comparison of expression profiles of isoforms of the CLDN18 gene, as detected with k4neo and STAR in the same set of GTEx samples. **(C)** Correlation of k-mer coverage along full-length transcript (k-mer fraction) and gene expression strength in TPM. Each dot depicts the TPM and k mer fraction of a CTA in an individual melanoma sample. **(D)** The sample rate by k4neo (top) and in expression databases (bottom) of 8 full-length cancer testis antigens per TCGA tumor cohorts (left) and healthy tissues (right). k4neo recovers tissue- and disease-specific expression profiles of CTAs.

Together, k4neo combines efficient k-mer indexing with a large, curated resource of healthy tissue and tumor samples to enable rapid, scalable tumor specificity assessment and antigen prioritization.

### k4neo captures tissue-specific expression profiles of isoforms and full-length cancer testis antigens

Next, we assessed whether k4neo can distinguish expression profiles of known isoforms. CLDN18 encodes a tight junction protein with two major isoforms that exhibit strong tissue specificity: CLDN18.1 is expressed in lung, stomach, and testis, whereas CLDN18.2 is expressed only in stomach and is a therapeutic target in several solid tumors^59,95,96^. Using the first splice junction of each isoform as representative sequences, k4neo accurately recapitulated their expected expression patterns across the healthy tissues (Fig. 3B). CLDN18.1 was detected in 96.55% of lung and 60% of stomach samples, whereas CLDN18.2 was detected predominantly in stomach (93.33%) and not in lung. These results were consistent with read-based RNA-seq quantification in GTEx (Pearson correlation (R² = 0.998, p < 3.9 × 10^−49^). Fig. 3B; Supplementary Fig. 3A) and matched the known expression profile of these isoforms. Indicating that k4neo can reproduce known biological findings.

Next, we tested if k4neo accurately recovers the tumor-associated expression profiles of known cancer testis antigens (CTA)^97,98^. Although k4neo technically provides only the presence or absence of k-mers across indexed samples, we investigated whether the fraction of detected k-mers for a full-length transcript can approximate quantitative gene expression measures, such as transcripts per million (TPM). When analyzing 18 CTAs in 73 melanoma samples^60,61^, we observed that the fraction of k-mers detected along the transcript sequence (k-mer ratio) correlated strongly with expression strength (R = 0.82, p < 2.2×10⁻¹; Fig. 3C; Supplementary Fig. S3B). We found an optimal threshold of 70% k-mer rate of full-length transcripts to detect expression > 2 TPM and k4neo achieved 96% accuracy in classifying genes as expressed (Supplementary Fig. S3B). Finally, we assessed whether k4neo captures the distinct expression profiles of CTAs across all tumor types and healthy tissues in the k4neo index. Compared to standard mapping-derived transcript expression profiles of the antigens, k4neo reproduced the expected expression profiles (Pearson correlation R² = 0.87, p < 3.2 × 10^-286^), characterized by broad expression in specific tumor types, high expression in testis tissue, and rare expression in healthy tissues (Fig. 3D).

### k4neo accurately detects tumor specificity across neoantigen classes

After evaluating non-mutated TAAs, we next assessed whether k4neo can accurately predict the tumor-specificity of neoantigen candidates derived from somatic mutations. First, we investigated germline single nucleotide polymorphisms (SNPs) and somatic single nucleotide variants (SNVs) obtained from whole-exome sequencing of 73 melanoma patients^60,61^. Using k4neo, we defined variants as tumor-specific, if the mutated transcript sequences were detected in less than 1% of all healthy tissue samples. As expected, germline SNPs were frequently detected in healthy tissues (89.5%), whereas somatic SNVs were largely absent (7.2%) and thereby k4neo classified 92.8% of somatic SNVs as tumor-specific (Fig. 4A). Notably, the k4neo sample rate for germline SNPs correlated with overall population allele frequency reported by the GnomAD database^99^ (R² = 0.56, p < 2.2 × 10⁻¹, Supplementary Fig. S4). Using the germline and somatic mutated sequences, we found an optimal sample rate threshold of 0.7%, at which k4neo achieved an overall classification accuracy of 91% and ROC-AUC of 95.3% in classifying somatic variants as tumor-specific and germline variants as not tumor-specific (Fig. 4B).

**Figure 4.**
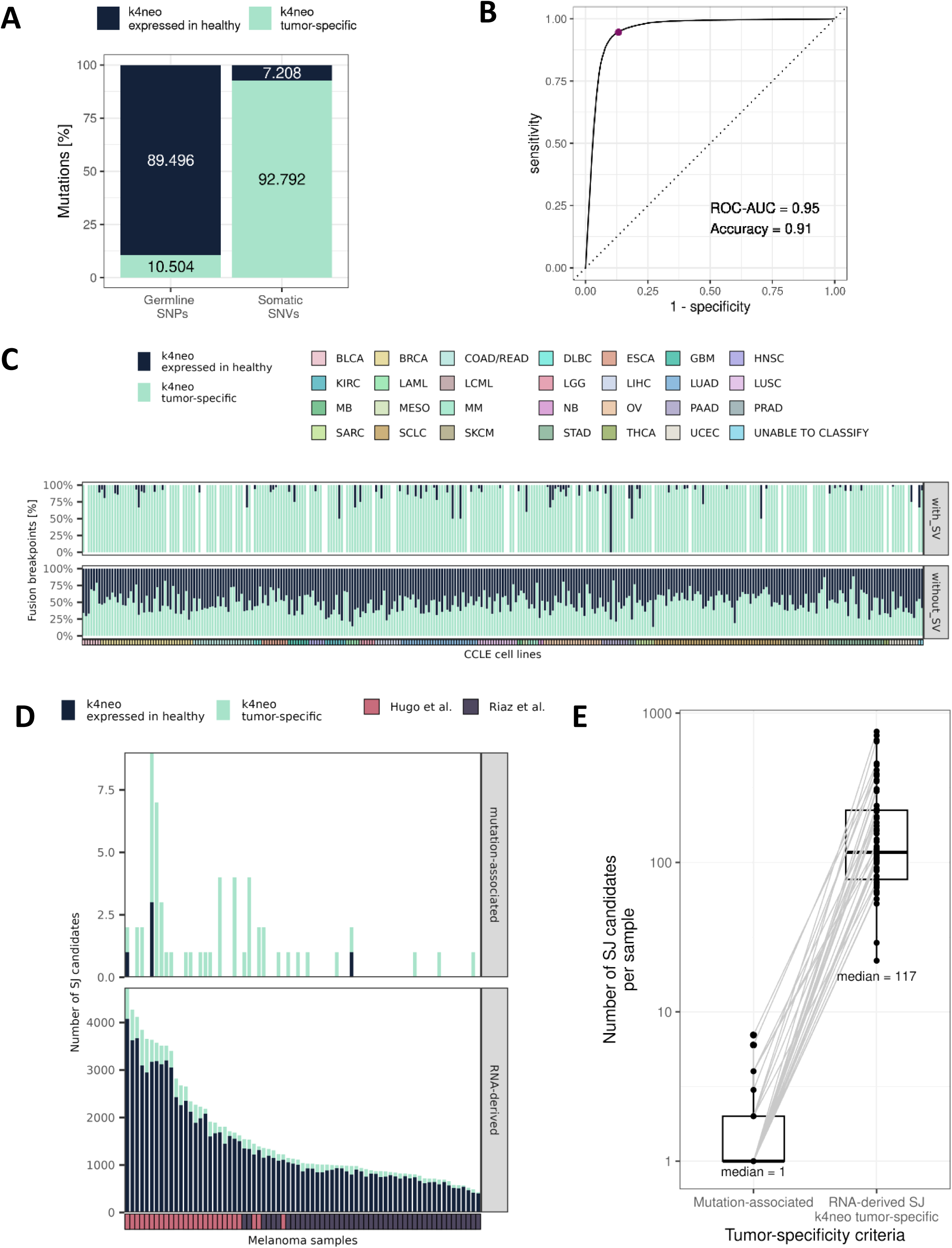
k4neo reliably detects tumor-specificity of neoantigen classes with genomic support for somatic origin. **(A)** The fraction of germline and somatic mutations in melanoma samples that were detected by k4neo in healthy tissues (more than 1% of healthy tissue samples) or not detected. **(B)** Receiver operating characteristic (ROC) curve for classifying somatic SNVs as tumor-specific and germline SNPs as not tumor-specific, by k4neo based on varying detection rates in healthy tissues. **(C)** Percent of gene fusions detected as tumor-specific or not tumor-specific by k4neo for gene fusions associated to structural variants from whole genome sequencing or gene fusion transcripts only detected from RNA-seq **(D)** Splice junctions per melanoma sample that are induced by somatic mutations (top) or detected only from RNA-seq (bottom). **(E)** Number of RNA-seq derived splice junctions candidates per melanoma sample that are mutation associated or predicted with k4neo to be tumor-specific.

Next, we assessed whether k4neo can predict the tumor specificity of gene fusion transcripts to support their use as neoantigens. Gene fusions are often identified from tumor RNA-seq data, and without matched whole-genome sequencing (WGS) data on structural variants (SVs), their tumor specificity remains uncertain. Using 327 tumor cell line samples with matched RNA-seq and WGS-based SV data, we compared k4neo-derived tumor specificity between fusion transcripts associated with somatic SV breakpoints and those supported only by RNA-seq. Fusion transcripts identified solely at the RNA level were frequently detected in healthy tissues (50.5%), indicating lower tumor specificity. In contrast, among SV-supported fusion transcripts, only 4.5% were detected in healthy tissues and 95.5% were classified as tumor-specific (Fig. 4C).

Analogously to SVs and gene fusions, somatic mutations can disrupt canonical splice motifs or create novel splice sites, giving rise to highly tumor-specific splice junctions that may encode neoantigens^18,19,100^. In a previous study of melanoma samples^48^, we reported an average of 1.7 mutation-associated splice junctions per patient and between 641 and 5,606 novel splice junctions without detectable mutational support. Using k4neo, we assessed the tumor specificity of splice junctions with and without somatic mutation support (Fig. 4D). Among RNA-seq–derived junctions lacking mutational support, 82% were detected in healthy tissues, indicating limited tumor specificity. In contrast, 91.5% of mutation-associated splice junctions were classified as tumor-specific by k4neo. In addition, we used the non-mutated junctions to quantify how many further tumor-specific junctions k4neo could recover to expand the antigen pool. k4neo identified a median of 117 tumor-specific non-mutated junctions per patient, compared with a median of 1 mutation-associated junction, substantially enlarging the candidate pool (Fig. 4E).

Together, these results demonstrate that k4neo accurately predicts tumor specificity of variant transcript sequences arising from somatic alterations, including mutations, gene fusions, and splice junctions, supporting its potential for prioritizing other antigen candidates.

### k4neo reveals highly tumor-specific splice junctions as individual and shared tumor antigen targets

Motivated by k4neo’s strong performance in predicting tumor specificity, we next sought to expand the antigen repertoire by identifying highly tumor-specific splice junctions in multiple tumor types. To this end, we developed NeoRasp, a novel end-to-end workflow that detects splice junctions from tumor RNA-seq, annotates their peptide and predicted MHC epitopes, and integrates k4neo for tumor-specificity filtering (Supplementary Fig. S5A). NeoRasp is implemented as a reproducible, containerized Snakemake pipeline and is released as open-source software with this study. We applied NeoRasp to nine cancer cohorts comprising 1,244 samples (Supplementary Table S7): glioblastoma (GBM, n=126)^101,102^, skin melanoma (SKCM, n=38)^60^, uveal melanoma (UVM, n=19)^87^, acute myeloid leukemia (AML, n=409)^103^, non-small cell lung cancer (NSCLC, n=199)^104^, small cell lung cancer (SCLC, n=57)^105^, triple negative breast cancers (BRCA/TNBC, n=57)^16^, colorectal cancers (CRC, n=88)^106^ and sarcomas (SARC, n=147)^107^. After requiring sufficient read support, excluding junctions matching annotated GENCODE transcripts, and removing junctions present in a long-read–based healthy-tissue exclusion set^108,109^, we identified a median of 884 novel candidate junctions per UVM sample (Fig. 5A). Filtering these candidates with k4neo for tumor specificity reduced this number by more than tenfold, to a median of 79 tumor-specific junctions per UVM sample. Across tumor types, we detected a median of 4–80 tumor-specific junctions per sample (Fig. 5B). Applying the same pipeline to independent healthy-tissue cohorts from blood^81^, brain^82^, heart^83^ and liver^84^ (n=248), which were not included in the healthy-tissue index, yielded a median of zero tumor-specific junctions, consistent with a low false-positive rate of k4neo (Fig. 5B).

**Figure 5.**
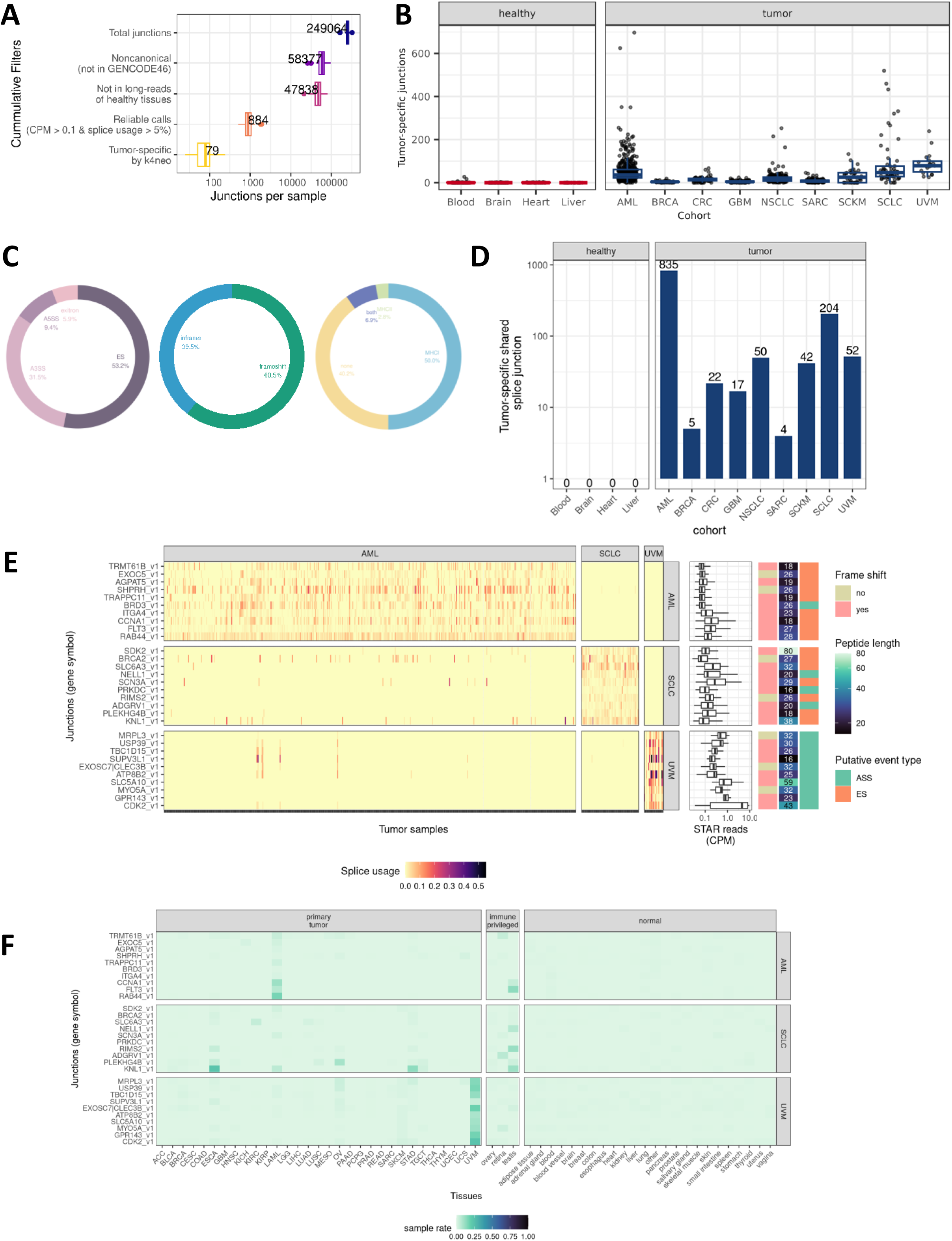
The recurrence of shared tumor-specific splice junctions. **(A)** Cumulatively applied filtering steps in the NeoRasp splice junction pipeline including k4neo tumor-specificity filtering (y-axis) and number of resulting splice-junctions per sample (x-axis) for 19 UVM samples. **(B)** Detected novel tumor-specific junctions per sample in healthy (negative control) and tumor cohorts. **(C)** Fraction of alternative splicing event types (left), their impact on reading frame (in-frame or frameshift, middle), and predicted MHC class I or class II binding epitopes (right). **(D)** Number of tumor-specific shared splice variants for tumor and healthy tissue negative control cohorts, whereby shared is defined as detection in at least 5% of samples per cohort. **(E)** Top 10 most recurrent splice junctions with annotated peptide change for AML, SCLC and UVM cohorts (y-axis). Splice junction usage across tumor samples (x-axis) is shown in left heatmap, alongside junction read support across detecting samples (middle, horizontal boxplots), frame shift annotation, novel peptide length and splice junction event type. **(G)** Recurrence of candidates in healthy tissues and confirmation in unmatched TCGA tumor cohorts using k4neo).

Next, we examined the tumor-specific splice junctions identified in individual patients as source of tumor-specific T-cell antigen candidates. After restricting to junctions for which an altered protein sequence could be annotated, we identified 4,772 candidates (Supplementary Fig. S5B). These comprised exon skipping (53.2%), alternative 3′ splice-site usage (31.5%), alternative 5′ splice-site usage (9.4%) and exitrons (5.9%). Overall, 60.5% of candidates generated frameshift peptides, whereas 39.5% preserved the reading frame. Using patient-specific HLA alleles for MHC binding prediction, we found that 59.7% of peptides contained at least one epitope predicted to be presented by MHC class I and/or class II (Fig. 5C).

We next assessed whether these highly tumor-specific splice junctions were recurrently detected in patients of the same tumor type and potentially qualify as shared antigen candidates. While no recurrent junctions were detected in the healthy control cohorts, between 4 and 835 shared tumor-specific junctions were identified per tumor type (Fig. 5D). After restricting to peptide altering events, we obtained few or no shared candidates for GBM (n=1), TNBC (n=2), CRC (n=4), NSCLC (n=0), and SARC (n=0), but between 33 and 109 shared candidates for UVM (n=33), SCLC (n=73), and AML (n=109), which were detected in up to 80% of patients (Supplementary Fig. S5B, C). We next focused on the 10 most recurrent junctions in AML, UVM and SCLC (Fig. 5F). These junctions clustered strongly by tumor type, while some were expressed across entities. Exon-skipping events predominated in AML and SCLC, whereas alternative splice-site usage prevailed in UVM. Most junctions introduced frameshifts and encoded novel peptide sequences up to 80 amino acids in length (Fig. 5E). Next, we assessed recurrence in the larger TCGA cohorts. We queried these junction sequences with k4neo against the TCGA index and confirmed recurrent detection of the AML and UVM junctions at high rates in the corresponding TCGA tumor samples (Fig. 5F).

Using patient-specific HLA types for MHC epitope binding prediction, we identified recurrent junctions that are expressed and generated at least one predicted strong MHC class I or class II binder in more than 25% of patients (AML n=7, SCLC n=8, UVM n=9; Fig. 6A). We then asked how many shared variants would need to be combined to provide broader patient coverage. In SCLC, combining four exon-skipping events (KNL1, PLEKHG4B, ADGRV1 and SCN3A) covered 98.3% of patients with at least one predicted strong-binding epitope (Fig. 6B). Adding splice variants in SDK2 and SLC6A3 increased coverage to 94.9% of patients with at least two predicted binders. In AML, the top-ranked candidates covered 83.9% of patients with at least one epitope, 63.3% with at least two epitopes and 45.7% with three or more epitopes (Fig. 6C). In UVM, all top recurrent junctions were predicted to encode strong-binding epitopes in expressing samples (Fig. 6D).

**Figure 6.**
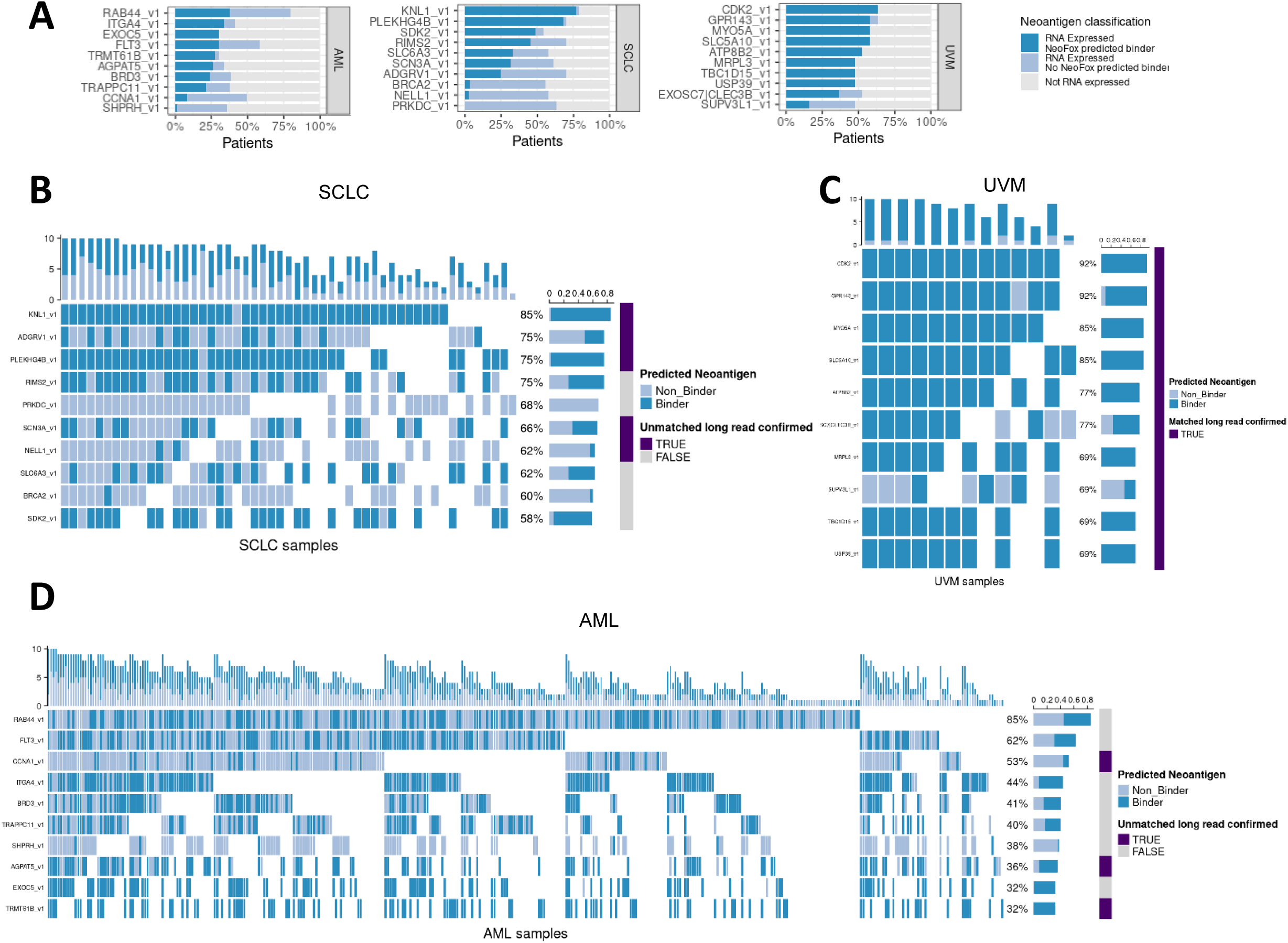
The shared antigen landscape from tumor-specific splice junctions in SCLC, UVM and AML. **(A)** For the 10 most recurrent splice junctions-derived antigen candidates per tumor cohort, the patient coverage is shown as percent of patients in tumor cohorts expressing the splice junction (light blue) and percent of patients with predicted MHC epitope using patient-specific HLA types (dark blue). **(B, C, D)** The top 10 recurrent candidate splice junctions (y-axis) in SCLC **(B)**, AML **(C)**, and SCLC **(D)** are shown for each expressing sample (x-axis) and are represented as MHC class I or II binder or non-binder (blue/gray). For each candidate junction it is indicated whether it could be confirmed in patient- or tumor-type-matched long-read sequencing data. Bars on the right side (dark purple) indicate confirmation with RNA long-read sequencing.

Finally, we confirmed the 10 most recurrent tumor-specific splice junctions using sample-matched or indication-matched long-read RNA sequencing. All top recurrent UVM candidates were confirmed in sample-matched long-read data (Fig. 6C), and six SCLC and three AML candidates were confirmed in indication-matched long-read datasets (Fig. 6B,D), indicating a high rate of independent full-length transcript validation. Notably, a multi-exon-skipping transcript variant of AGPAT5 (AGPAT5_v1) was expressed at higher levels than the canonical reference transcript in the long-read data. AGPAT5_v1 introduces a frameshift and encodes 19 novel amino acids; it was recurrently detected in 36% of AML samples, of which more than 80% were predicted to present MHC epitopes derived from this variant.

Together, these results show that stringent tumor-specificity assessment with k4neo enables the identification of both patient-specific and recurrent tumor-specific splice junctions. The recurrent junctions are broadly expressed, can be confirmed by long-read sequencing, and encode predicted MHC-binding epitopes that cover a substantial fraction of patients, highlighting their potential as shared tumor-specific antigens for immunotherapeutic applications.

## Discussion

Expanding the tumor antigen repertoire beyond somatic SNVs is essential for extending personalized immunotherapies, including neoantigen vaccines targeting low-TMB malignancies. To date, the use of non-canonical transcript variants as antigen candidates has been hindered by the lack of efficient, accurate methods to confirm their tumor specificity^3,30,110,111^. To address this, we developed k4neo, which combines pre-built k-mer indices of 18,960 healthy (GTEx, SRA), 10,328 tumor (TCGA) and 737 adjacent normals (TCGA) RNA-seq samples with curated metadata to annotate any input transcript sequence with expression profiles across diverse human tissues and tumor types and to accurately predict its tumor specificity. Unlike existing approaches^34,35^, k4neo is mapping-free and transcript-class agnostic: it scores splice junctions, gene fusions, full-length isoforms, or point mutations within a single framework, without a precomputed event-specific exclusion list. Its computational efficiency enables per-patient screening, allowing thousands of candidate sequences of any class to be queried against thousands of unmatched healthy samples. Because tumor-specificity is a shared prerequisite, the same screening applies not only to neoantigen vaccines but also to the selection of antibody-, bispecific-, and CAR-/TCR-T-cell targets, where on-target/off-tumor toxicity is a central safety concern^15,34^.

Our benchmark of nine k-mer indexing modes showed that, while detection accuracy on short splice-junction queries was high across all tested methods, computational scalability differed substantially. Several tools exhibited exponential growth in build time or memory consumption with cohort size, and we could not reproduce the reported scalability of MetaGraph, observing prohibitive memory use consistent with previous findings^44^. Raptor in HIBF mode offered the best overall trade-off, ranked among the most accurate methods, with the fastest builds, smallest indices, and sub-minute queries, and was therefore selected as k4neo’s default backend.

k4neo robustly recapitulated established tissue-specific isoform expression patterns, exemplified by CLDN18, as well as the tumor-associated expression of cancer-testis antigens. Moreover, it accurately predicted tumor specificity across three classes of somatic alteration-derived neoantigens antigens. Importantly, somatic SNVs, SV-supported fusions, and mutation-associated splice junctions provide rare cases for which tumor-specificity can be established independently from genomic evidence. k4neo’s high accuracy on these benchmarks therefore qualifies its use on the far larger universe of non-canonical transcript variants, e.g. RNA-only fusions, novel splice junctions, retained introns, transposons, or cryptic ORFs, for which currently no orthogonal ground truth exists for tumor-specificity.

Applied to nine tumor cohorts, k4neo recovered a median of 4–80 tumor-specific splice junctions per patient, representing a more than tenfold reduction from raw candidates, while held-out healthy cohorts yielded a median of zero, consistent with a low false-positive rate. Across 4,772 protein-altering junctions, 59.7% encoded peptides predicted to bind patient-specific MHC alleles, substantially expanding the per-patient antigen pool. Among recurrent events, shared tumor-specific junctions were identified in AML, SCLC, and UVM and independently confirmed by long-read sequencing, including the multi-exon-skipping variant AGPAT5_v1, recurrent in 36% of AML samples and predicted to yield strong MHC binders in >80% of patients expressing the variant. The lower number of shared candidates in GBM, NSCLC, SARC, BRCA, CRC, and SKCM, despite reports of recurrent aberrant splicing in other studies^75,112^, likely reflects differences in detection pipelines, our stringent specificity threshold, and limited cohort size.

Despite its strong performance, also some limitations remain. First, tissue coverage in the healthy tissue index is uneven, particularly for rare or under-studied tissues such as thymus, which are represented by relatively few samples and thus may underestimate off-tumor expression. Consequently, this underscores the importance of extension of healthy reference tissue cohorts. Second, the optimal tumor-specificity threshold remains an open question: relaxing the ≤1% cut-off used here increases candidate yield exponentially (Supplementary Fig. S5E) but at unknown cost to immunogenicity or off-tumor toxicity, and the scarcity of experimentally validated non-canonical antigens currently limits empirical calibration of this trade-off. Third, k4neo currently operates on a binary presence/absence index. Although the fraction of detected k-mers along a transcript correlates with expression, per-sample quantitative measures, e.g. for prioritizing tumor-associated and shared antigens, would require quantitative k-mer data structures with sufficient computational scalability to large healthy tissue cohorts, which is an active area of development^39,45,46,113^. k4neo’s modular architecture supports drop-in replacement of its indexing backend through method-specific parsers, allowing it to incorporate such advances as they mature. In addition, the k4neo index can be incrementally expanded to more cohorts of healthy and tumor samples when available. k4neo assesses tumor specificity at the RNA level; for surface-directed modalities such as antibodies or CAR-T cells, its profiles provide an efficient first-line filter to prioritize candidates that should subsequently be confirmed at the protein and cell-surface level.

Long-read RNA sequencing not only resolves full-length transcript variants in tumors^87,114–116^, but also offers a complementary route to assessing tumor-specificity. Recent atlases have revealed thousands of additional full-length isoforms in healthy tissues^108,109^, refining the universe of reference transcripts. While long-read coverage across tissues and developmental stages remains too sparse to serve as a standalone filter, combining its sensitivity for resolving transcript complexity with the broad sample coverage of mapping-free k-mer indices from short-read data will further strengthen tumor-specificity assessment.

Together, k4neo turns the petabyte-scale corpus of public healthy RNA-seq into an accessible, searchable resource for assessing the tumor specificity of any candidate transcript sequence, bridging algorithmic advances in k-mer indexing with clinically relevant antigen discovery. Combined with patient-level RNA-seq pipelines, it enables both individualized antigen prioritization in low-TMB tumors and the discovery of recurrent, shared antigens for off-the-shelf vaccines, with potential applicability to additional therapeutic modalities such as antibody- and cell-based therapies. With its curated healthy-tissue reference atlas, k4neo thus provides a community resource that broadens the safe and actionable antigen repertoire for next-generation cancer immunotherapy.

## Declarations

## Author contributions

Conceptualization: JI. Methodology: JH, JI. Software: JH, JI, LK, ÖM. Data curation: JH, FL, DW, RK. Formal analysis: JH, JI, FL, LK. Supervision: DW, MS, US, JI. Visualization: JH, JI. Writing—original draft: JH, JI. Writing—reviewing and editing: JH, FL, JI, DW, BS, MS, MMG, US, JI. All authors read and approved the final manuscript.

## Data availability

Pre-built k-mer indices of 2,256 RNA-seq samples obtained from SRA, ENCODE and Gene expression Omnibus are available together with curated sample metadata via GitHub (https://github.com/TRON-Bioinformatics/k4neo). Sequencing data provided by the TCGA were obtained through dbGaP accession phs000178.v11.p8 and sequencing data from GTEx were obtained through dbGaP accession phs000424.v11.p2. We provide a WDL workflow to construct the k-mer indices of access-restricted GTEx and TCGA samples in the TERRA bioinformatics cloud on GitHub.

## Code availability

k4neo python package is available on GitHub (https://github.com/TRON-Bioinformatics/k4neo). Curated metadata for k4neo annotation and workflows for k-mer index construction are also available and documented in the GitHub repository. The NeoRasp pipeline for splice junction and antigen detection is available as Snakemake workflow on GitHub (https://github.com/TRON-Bioinformatics/NeoRasp).

## Conflict of interest

U.S. is co-founder, chief executive officer and stock owner of BioNTech SE. J.I. and J.H. are listed as inventors on a pending patent application related to this work (PCT/EP2025/063053). The remaining authors declare no competing interests.

## Acknowledgement

In the following manuscript, public sequencing data from multiple studies were used. We gratefully acknowledge the patients and their families for donating tissue samples, the study authors from the originating laboratories responsible for obtaining the tissue specimens, as well as the submitting laboratories where the sequence data were generated and shared via the European Nucleotide Archive, NCBI Sequencing Read archive, Gene Expression Omnibus and ENCODE. The authors used large language models to improve the language and readability of the manuscript. All scientific content was reviewed and verified by the authors. M.H.S. received funding through the Deutsche Forschungsgemeinschaft (DFG, German Research Foundation) under Germanýs Excellence Strategy – EXC 2026, Cardio-Pulmonary Institute, Project ID: 39064989.

---

**Supplementary Figure S1.**
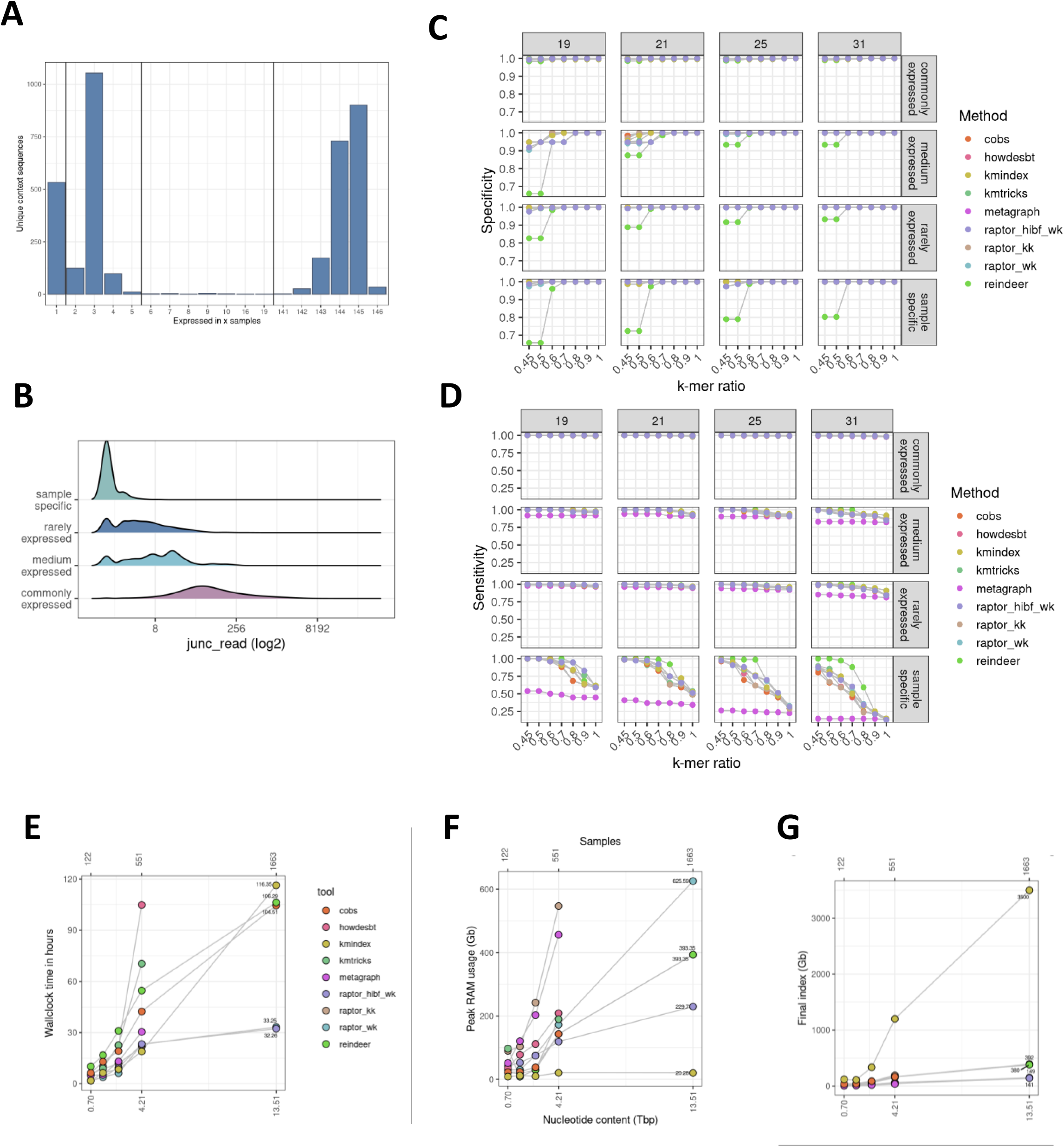
K-mer indexing methods benchmark data and results. **(A)** The Number of junction sequences per sample expressing the junction in the junction benchmark ground truth dataset. **(A)** Distribution of number of reads supporting the junctions from benchmark data by the four groups of expression breath across samples. **(C)** Specificity and **(D)** Sensitivity by method for multiple expression classes of splice junctions (rows) and different parameters for k-mer size (columns) and k-mer ratio thresholds (x-axis). **(E)** Computational efficiency of k-mer methods by cohort size as time to construct index by cohort size (x-axis, indicated as nucleotide content in Tbp at bottom and number of samples at top). **(F)** Peak memory usage in Gb during index construction by cohort size. **(G)** Final total file size of index in Gb by cohort size.

**Supplementary Figure S2.**
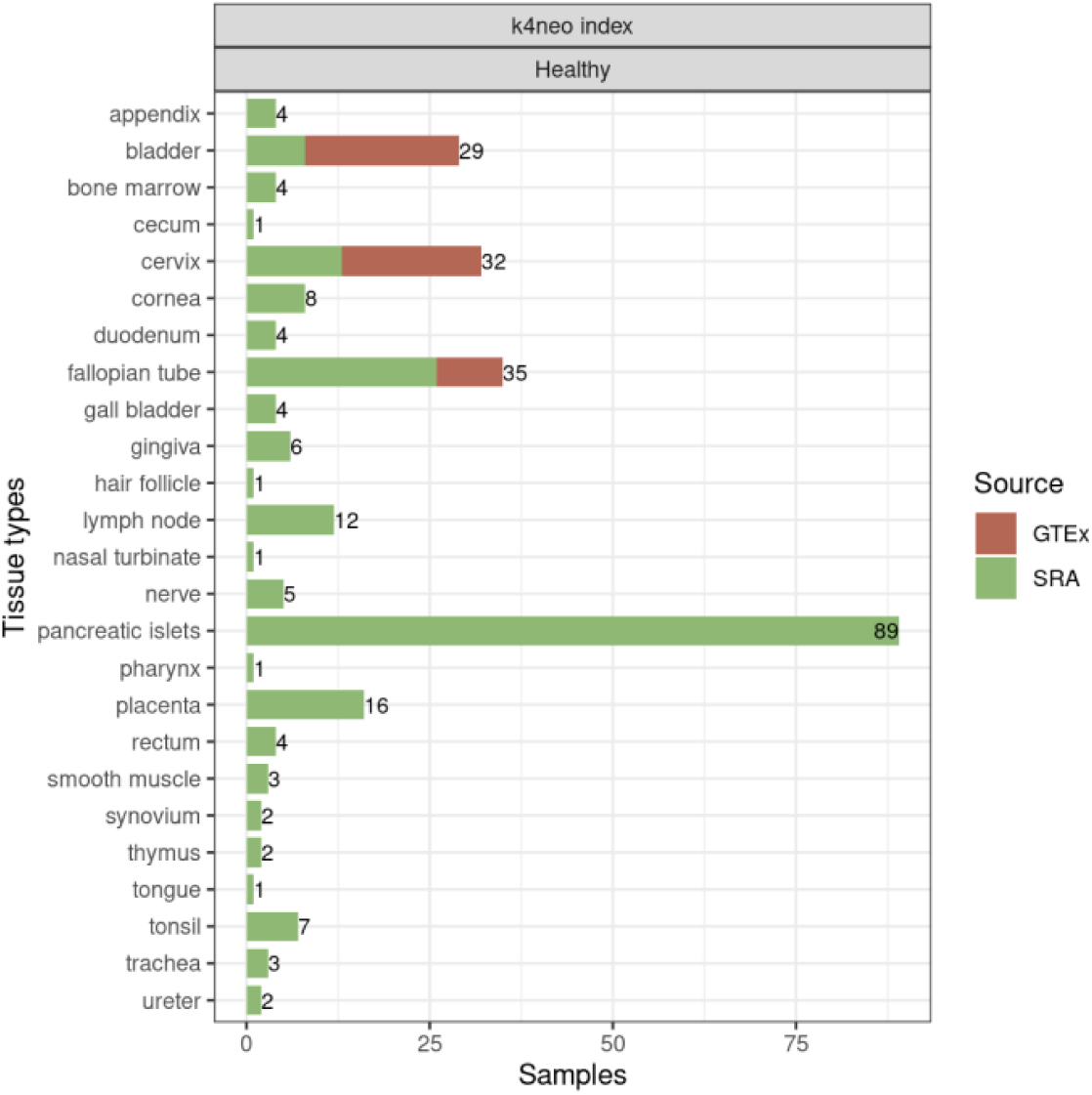
Overview of additional tissue samples in healthy tissue index: Additional tissue samples grouped as other in analysis are listed and annotated by origin.

**Supplementary Figure S3.**
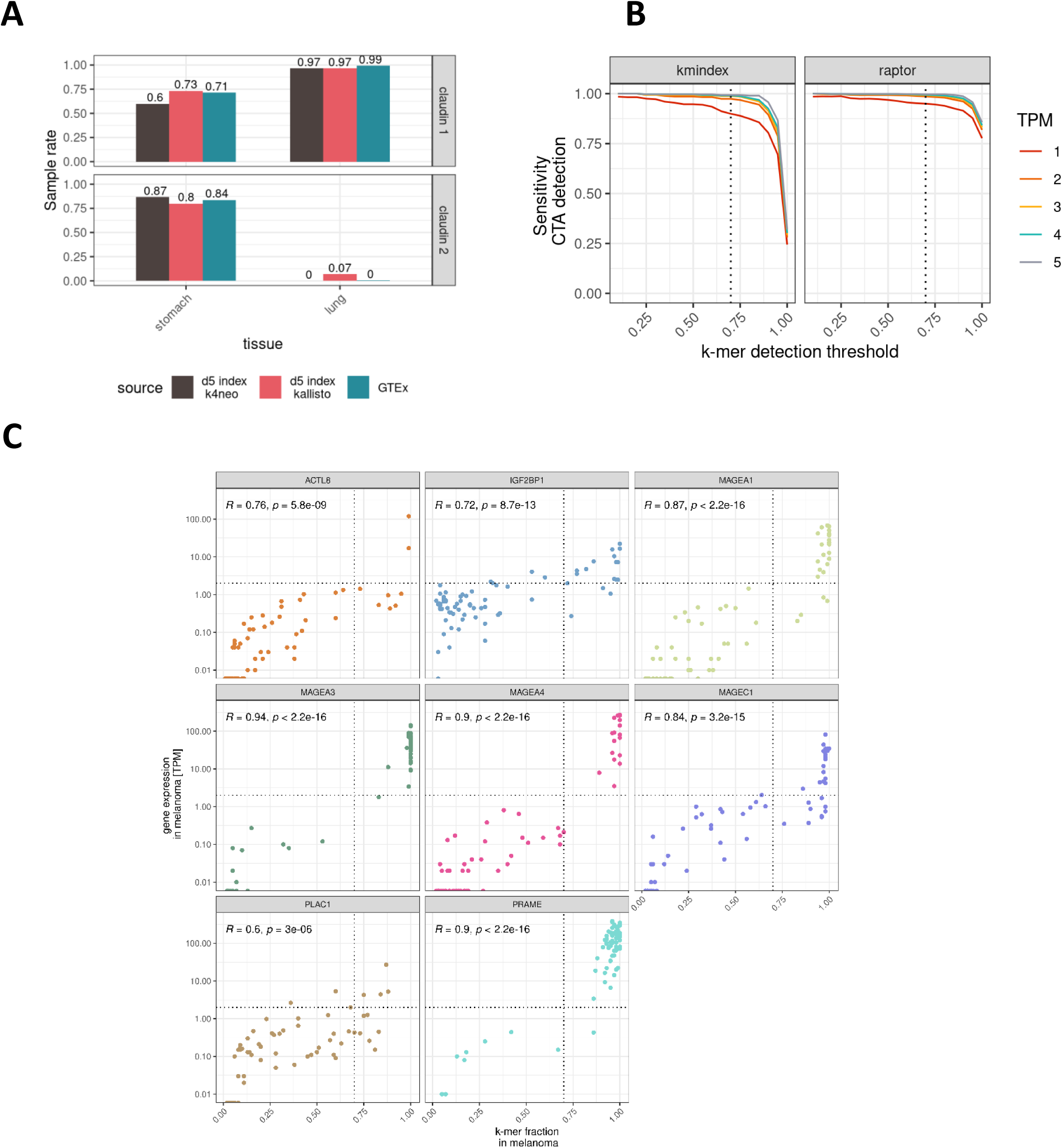
Full-length transcript detection sensitivity. **(A)** Comparison CLDN18 sample rates of k4neo (SRA index), kallisto (SRA lung & stomach samples) and GTEx STAR counts. **(B)** Detection sensitivity of kmindex and raptor to detect full-length CTA transcripts at varying TPM cutoffs (1 to 5) by k-mer ratio. **(C)** Correlation of k-mer ratio along the sequence and expression value in TPM. Each dot depicts a melanoma sample.

**Supplementary Figure S4.**
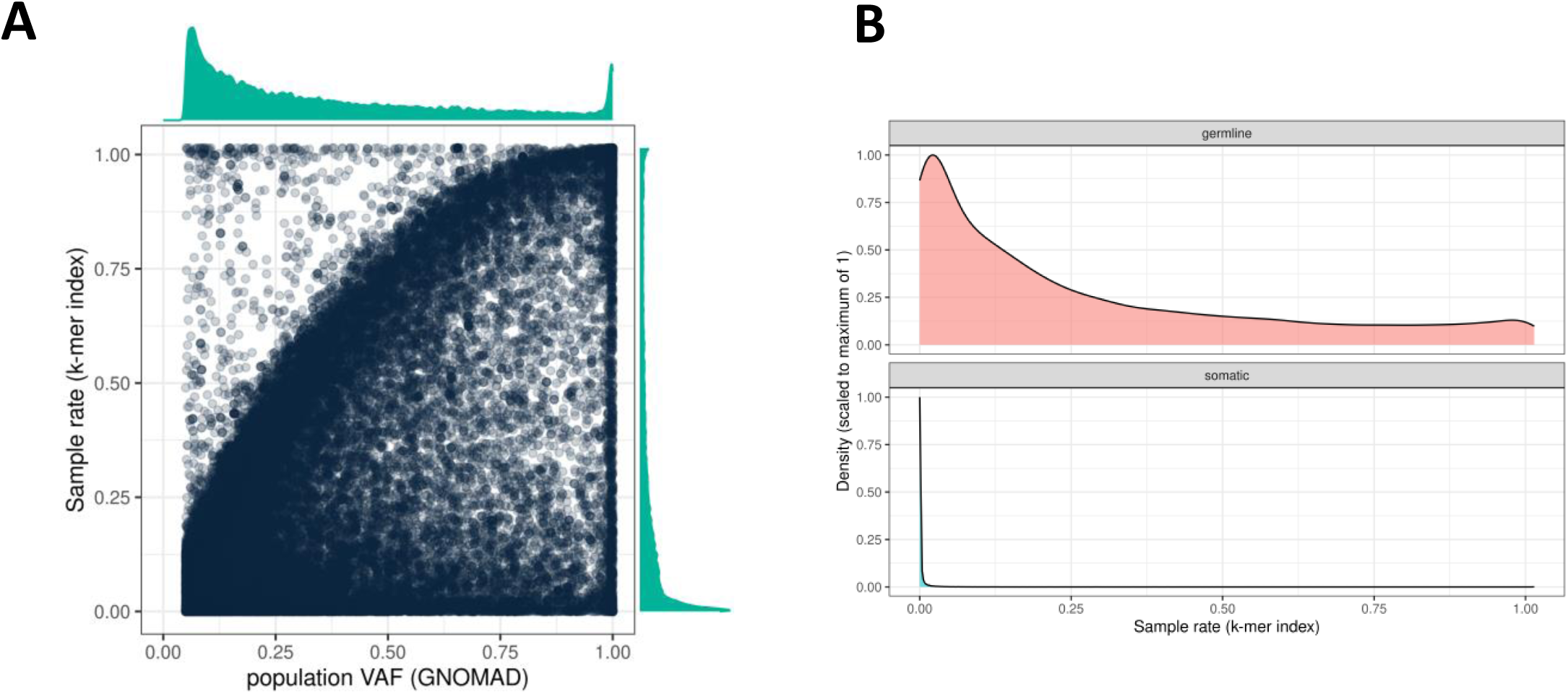
Sample rate of germline and somatic point mutations by k4neo. **(A)** Sample rate in healthy tissue index by k4neo (y-axis) by population variant frequency in gnomAD (x-axis) for germline variants in melanoma samples. **(B)** Distribution of sample rate in healthy tissue index by k4neo for germline variants (top) and somatic SNVs (bottom) for melanoma samples.

**Supplementary Figure S5.**
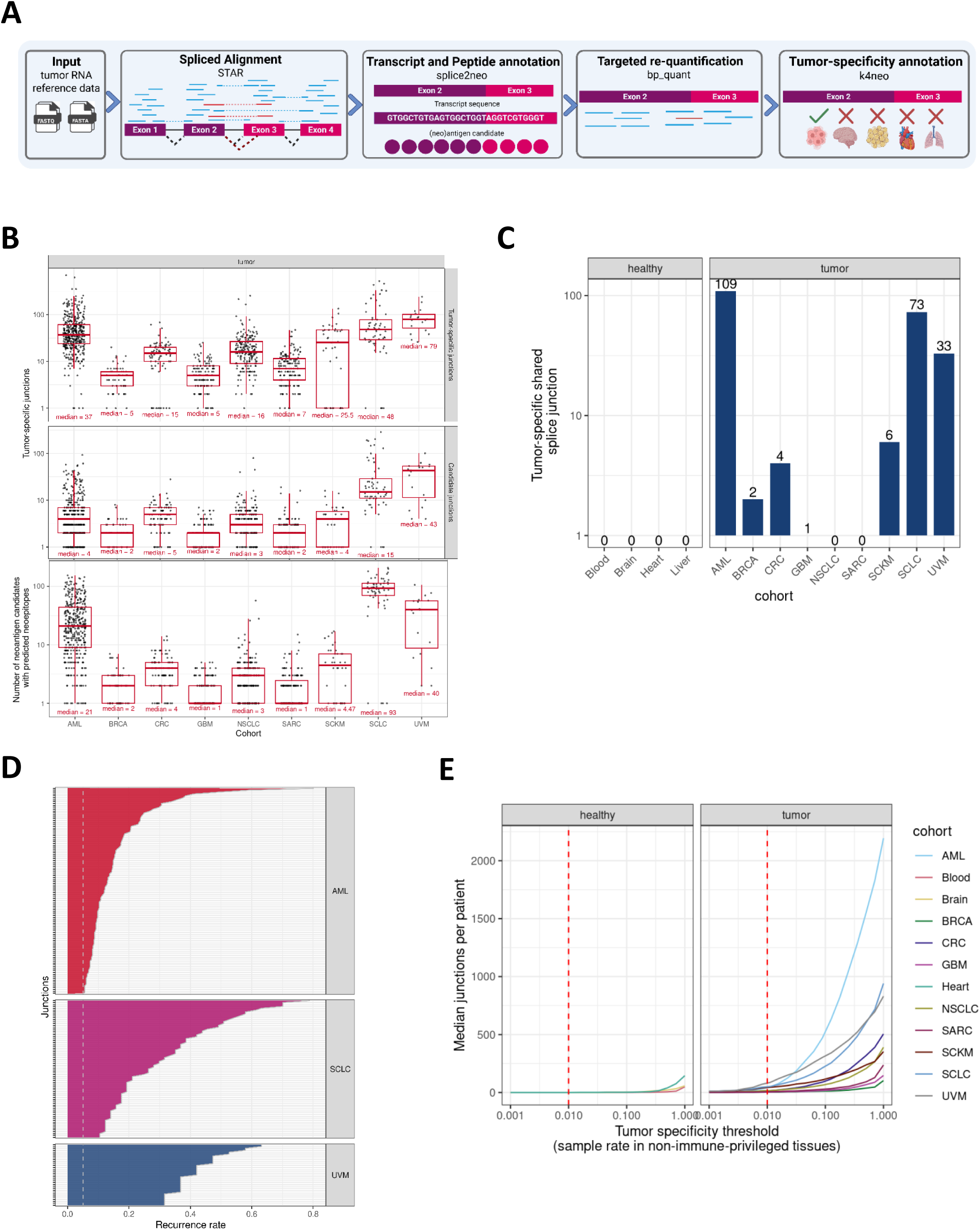
Detection of tumor-specific splice variants. **(A)** Overview of the NeoRasp pipeline to detect splice junctions from RNA-seq and k4neo for tumor-specificity detection. **(B)** Number of tumor-specific and peptide altering splicing junctions (candidates) in normal and tumor cohorts. **(C)** Number of tumor-specific shared splice variants affecting the coding sequence for tumor cohort and negative control healthy tissue cohorts. **(D)** Recurrence rate (x-axis) of tumor-specific shared splice junction candidates (y-axis) in AML, SCLC and UVM cohorts. Vertical line depicts recurrence criterion of 5%. **(E)** Median number of tumor-specific splice junction per sample (y-axis) by varying tumor-specificity cutoff for detection rate in none-immune-privileged healthy tissue samples (x-axis) for negative control healthy tissues (left) and tumor cohorts (right).

## Notes

https://github.com/TRON-Bioinformatics/k4neo

### Competing Interest Statement

U.S. is co-founder, chief executive officer and stock owner of BioNTech SE. JH and JI are listed as inventors on a pending patent application related to this work (PCT/EP2025/063053). The remaining authors declare no competing interests.

## References

1. Gilboa, E. The Makings of a Tumor Rejection Antigen. Immunity 11, 263–270 (1999).

2. Schumacher, T. N. & Schreiber, R. D. Neoantigens in cancer immunotherapy. Science 348, 69–74 (2015).

3. Lang, F., Schrörs, B., Löwer, M., Türeci, Ö. & Sahin, U. Identification of neoantigens for individualized therapeutic cancer vaccines. Nat. Rev. Drug Discov. 1–22 (2022) doi:10.1038/s41573-021-00387-y.

4. Sahin, U. et al. Personalized RNA mutanome vaccines mobilize poly-specific therapeutic immunity against cancer. Nature 547, 222–226 (2017).

5. Snyder, A. et al. Genetic Basis for Clinical Response to CTLA-4 Blockade in Melanoma. N. Engl. J. Med. 371, 2189–2199 (2014).

6. Klempner, S. J. et al. Tumor Mutational Burden as a Predictive Biomarker for Response to Immune Checkpoint Inhibitors: A Review of Current Evidence. The Oncologist 25, e147–e159 (2020).

7. Ott, P. A. et al. An immunogenic personal neoantigen vaccine for patients with melanoma. Nature 547, 217–221 (2017).

8. Hilf, N. et al. Actively personalized vaccination trial for newly diagnosed glioblastoma. Nature 1 (2018) doi:10.1038/s41586-018-0810-y.

9. Sahin, U. & Türeci, Ö. Personalized vaccines for cancer immunotherapy. Science 359, 1355–1360 (2018).

10. Blass, E. & Ott, P. A. Advances in the development of personalized neoantigen-based therapeutic cancer vaccines. Nat. Rev. Clin. Oncol. 1–15 (2021) doi:10.1038/s41571-020-00460-2.

11. Kandoth, C. et al. Mutational landscape and significance across 12 major cancer types. Nature 502, 333–339 (2013).

12. Sha, D. et al. Tumor Mutational Burden as a Predictive Biomarker in Solid Tumors. Cancer Discov. 10, 1808–1825 (2020).

13. Rojas, L. A. et al. Personalized RNA neoantigen vaccines stimulate T cells in pancreatic cancer. Nature 1–7 (2023) doi:10.1038/s41586-023-06063-y.

14. Keskin, D. B. et al. Neoantigen vaccine generates intratumoral T cell responses in phase Ib glioblastoma trial. Nature 565, 234–239 (2019).

15. Jafry, R. Z. et al. Efficacy and safety of approved cellular therapies and bispecific antibodies in solid tumors: the current state. *Discov*. Oncol. 17, 637 (2026).

16. Weber, D. et al. Accurate detection of tumor-specific gene fusions reveals strongly immunogenic personal neo-antigens. Nat. Biotechnol. 40, 1276–1284 (2022).

17. Yang, W. et al. Immunogenic neoantigens derived from gene fusions stimulate T cell responses. Nat. Med. 1 (2019) doi:10.1038/s41591-019-0434-2.

18. Lang, F., et al. Prediction of tumor-specific splicing from somatic mutations as a source of neoantigen candidates. Bioinforma. Adv. 4, vbae080 (2024).

19. Kahles, A. et al. Comprehensive Analysis of Alternative Splicing Across Tumors from 8,705 Patients. Cancer Cell 0, (2018).

20. Jayasinghe, R. G. et al. Systematic Analysis of Splice-Site-Creating Mutations in Cancer. Cell Rep. 23, 270–281.e3 (2018).

21. Smart, A. C. et al. Intron retention is a source of neoepitopes in cancer. Nat. Biotechnol. 10.1038/nbt.4239 (2018) doi:10.1038/nbt.4239.

22. Wang, T.-Y. et al. A pan-cancer transcriptome analysis of exitron splicing identifies novel cancer driver genes and neoepitopes. Mol. Cell 81, 2246–2260.e12 (2021).

23. Ott, P. A. The promises and challenges of neoantigen cancer vaccines. Nat. Biotechnol. 44, 740–751 (2026).

24. Anilkumar, A. S., Thomas, S. M. & Veerabathiran, R. Next-generation cancer vaccines: targeting cryptic and non-canonical antigens for precision immunotherapy. Explor. Target. Anti-Tumor Ther. 6, 1002338 (2025).

25. Ely, Z. A. et al. Pancreatic cancer–restricted cryptic antigens are targets for T cell recognition. Science 388, eadk3487 (2025).

26. Attig, J. et al. LTR retroelement expansion of the human cancer transcriptome and immunopeptidome revealed by de novo transcript assembly. Genome Res. 29, 1578–1590 (2019).

27. Laumont, C. M. et al. Noncoding regions are the main source of targetable tumor-specific antigens. Sci. Transl. Med. 10, eaau5516 (2018).

28. Bonté, P.-E. et al. Single-cell RNA-seq-based proteogenomics identifies glioblastoma-specific transposable elements encoding HLA-I-presented peptides. Cell Rep. 39, 110916 (2022).

29. Latysheva, N. S. & Babu, M. M. Discovering and understanding oncogenic gene fusions through data intensive computational approaches. Nucleic Acids Res. 44, 4487–4503 (2016).

30. Rwandamuriye, F. X., Redwood, A. J., Creaney, J. & Robinson, B. W. S. Hidden Targets in Cancer Immunotherapy: The Potential of “Dark Matter” Neoantigens. Vaccines 14, (2026).

31. Lonsdale, J. et al. The Genotype-Tissue Expression (GTEx) project. Nat. Genet. 45, 580–585 (2013).

32. Weinstein, J. N. et al. The Cancer Genome Atlas Pan-Cancer analysis project. Nat. Genet. 45, 1113–1120 (2013).

33. EMBL-EBI. ENA statistics – reads growth - reads doubling time. (2023).

34. Li, G., Schnell, D., Bhattacharjee, A., Yarmarkovich, M. & Salomonis, N. Quantifying tumor specificity using Bayesian probabilistic modeling for drug and immunotherapeutic target discovery. *Cell Rep*. Methods 4, 100900 (2024).

35. Wilks, C. et al. recount3: Summaries and queries for large-scale RNA-seq expression and splicing. Genome Biol. 22, 323 (2021).

36. Weber, D. et al. Accurate detection of tumor-specific gene fusions reveals strongly immunogenic personal neo-antigens. Nat. Biotechnol. 1–9 (2022) doi:10.1038/s41587-022-01247-9.

37. Bray, N. L., Pimentel, H., Melsted, P. & Pachter, L. Near-optimal probabilistic RNA-seq quantification. Nat. Biotechnol. 34, 525–527 (2016).

38. Patro, R., Duggal, G., Love, M. I., Irizarry, R. A. & Kingsford, C. Salmon provides fast and bias-aware quantification of transcript expression. Nat. Methods 14, 417–419 (2017).

39. Marchet, C., et al. Data structures based on k-mers for querying large collections of sequencing data sets. Genome Res. 10.1101/gr.260604.119 (2020) doi:10.1101/gr.260604.119.

40. Lemane, T., Medvedev, P., Chikhi, R. & Peterlongo, P. kmtricks: Efficient and flexible construction of Bloom filters for large sequencing data collections. Bioinforma. Adv. 2, vbac029 (2022).

41. Harris, R. S. & Medvedev, P. Improved representation of sequence bloom trees. Bioinforma. Oxf. Engl. 36, 721–727 (2020).

42. Bingmann, T., Bradley, P., Gauger, F. & Iqbal, Z. COBS: A Compact Bit-Sliced Signature Index. ArXiv Databases (2019).

43. Mehringer, S. et al. Hierarchical Interleaved Bloom Filter: enabling ultrafast, approximate sequence queries. Genome Biol. 24, 131 (2023).

44. Lemane, T. et al. Indexing and real-time user-friendly queries in terabyte-sized complex genomic datasets with kmindex and ORA. Nat. Comput. Sci. 4, 104–109 (2024).

45. Karasikov, M. et al. MetaGraph: Indexing and Analysing Nucleotide Archives at Petabase-scale. bioRxiv 2020.10.01.322164 (2020) doi:10.1101/2020.10.01.322164.

46. Marchet, C., Iqbal, Z., Gautheret, D., Salson, M. & Chikhi, R. REINDEER: Efficient indexing of k-mer presence and abundance in sequencing datasets. Bioinforma. Oxf. Engl. 36, i177–i185 (2020).

47. Dobin, A. et al. STAR: Ultrafast universal RNA-seq aligner. Bioinforma. Oxf. Engl. 29, 15–21 (2013).

48. Lang, F. et al. Prediction of tumor-specific splicing from somatic mutations as a source of neoantigen candidates. bioRxiv 10.1101/2023.06.27.546494 (2023) doi:10.1101/2023.06.27.546494.

49. Mohamadi, H., Khan, H. & Birol, I. ntCard: A streaming algorithm for cardinality estimation in genomics data. Bioinforma. Oxf. Engl. 33, 1324–1330 (2017).

50. Chikhi, R., Limasset, A. & Medvedev, P. Compacting de Bruijn graphs from sequencing data quickly and in low memory. Bioinforma. Oxf. Engl. 32, i201–i208 (2016).

51. Kokot, M., Dlugosz, M. & Deorowicz, S. KMC 3: Counting and manipulating k-mer statistics. Bioinforma. Oxf. Engl. 33, 2759–2761 (2017).

52. Tange, O. GNU Parallel - The Command-Line Power Tool. Login USENIX Mag. 36, 42–47 (2011).

53. Barrett, T. et al. NCBI GEO: Archive for functional genomics data sets–update. Nucleic Acids Res. 41, D991–5 (2013).

54. Luo, Y. et al. New developments on the Encyclopedia of DNA Elements (ENCODE) data portal. Nucleic Acids Res. 48, D882–D889 (2020).

55. Chen, S. Ultrafast one–pass FASTQ data preprocessing, quality control, and deduplication using fastp. iMeta 2, (2023).

56. Pandey, P. et al. Mantis: A Fast, Small, and Exact Large-Scale Sequence-Search Index. Cell Syst. 7, 201–207.e4 (2018).

57. Marchet, C. & Limasset, A. Scalable sequence database search using partitioned aggregated Bloom comb trees. Bioinforma. Oxf. Engl. 39, i252–i259 (2023).

58. Lemane, T. et al. kmindex and ORA: Indexing and real-time user-friendly queries in tera-bytes-sized complex genomic datasets. bioRxiv 10.1101/2023.05.31.543043 (2023) doi:10.1101/2023.05.31.543043.

59. Sahin, U. et al. Claudin-18 splice variant 2 is a pan-cancer target suitable for therapeutic antibody development. Clin. Cancer Res. Off. J. Am. Assoc. Cancer Res. 14, 7624–7634 (2008).

60. Hugo, W. et al. Genomic and Transcriptomic Features of Response to Anti-PD-1 Therapy in Metastatic Melanoma. Cell 165, 35–44 (2016).

61. Riaz, N. et al. Tumor and Microenvironment Evolution during Immunotherapy with Nivolumab. Cell 171, 934–949.e16 (2017).

62. Li, H. & Durbin, R. Fast and accurate short read alignment with Burrows-Wheeler transform. Bioinforma. Oxf. Engl. 25, 1754–1760 (2009).

63. Benjamin, D. et al. Calling Somatic SNVs and Indels with Mutect2. Preprint at 10.1101/861054 (2019).

64. Riesgo-Ferreiro, P. & Ritzel, C. TRON-Bioinformatics/tronflow-mutect2: Release v1.8.0. Zenodo 10.5281/zenodo.6592433 (2022).

65. Benjamin, D. et al. Calling Somatic SNVs and Indels with Mutect2. Preprint at 10.1101/861054 (2019).

66. Riesgo-Ferreiro, P. TRON-Bioinformatics/tronflow-haplotype-caller: v1.0.1. Zenodo 10.5281/zenodo.7313630 (2022).

67. Sherry, S. T. dbSNP: the NCBI database of genetic variation. Nucleic Acids Res. 29, 308–311 (2001).

68. The 1000 Genomes Project Consortium et al. A global reference for human genetic variation. Nature 526, 68–74 (2015).

69. The International HapMap Consortium. The International HapMap Project. Nature 426, 789–796 (2003).

70. Exome Aggregation Consortium et al. Analysis of protein-coding genetic variation in 60,706 humans. Nature 536, 285–291 (2016).

71. Ghandi, M. et al. Next-generation characterization of the Cancer Cell Line Encyclopedia. Nature 569, 503–508 (2019).

72. Lawrence, M., Gentleman, R. & Carey, V. rtracklayer: an R package for interfacing with genome browsers. Bioinformatics 25, 1841–1842 (2009).

73. Jaganathan, K. et al. Predicting splicing from primary sequence with deep learning. Cell 176, 535–548 (2019).

74. Cheng, J. et al. MMSplice: modular modeling improves the predictions of genetic variant effects on splicing. Genome Biol. 20, 1–15 (2019).

75. Kahles, A., Ong, C. S., Zhong, Y. & Rätsch, G. SplAdder: identification, quantification and testing of alternative splicing events from RNA-Seq data. Bioinformatics 32, 1840–1847 (2016).

76. Li, Y. I. et al. Annotation-free quantification of RNA splicing using LeafCutter. Nat. Genet. 50, 151–158 (2018).

77. Mölder, F. et al. Sustainable data analysis with Snakemake. F1000Research 10, 33 (2021).

78. Scheller, I. F., Lutz, K., Mertes, C., Yépez, V. A. & Gagneur, J. Improved detection of aberrant splicing with FRASER 2.0 and the intron Jaccard index. Am. J. Hum. Genet. 110, 2056–2067 (2023).

79. Langmead, B. & Salzberg, S. L. Fast gapped-read alignment with Bowtie 2. Nat. Methods 9, 357–359 (2012).

80. Lang, F., Riesgo-Ferreiro, P., Löwer, M., Sahin, U. & Schrörs, B. NeoFox: annotating neoantigen candidates with neoantigen features. Bioinformatics 37, 4246–4247 (2021).

81. Aguirre-Gamboa, R. et al. Deconvolution of bulk blood eQTL effects into immune cell subpopulations. BMC Bioinformatics 21, 243 (2020).

82. Ramaker, R. C. et al. Post-mortem molecular profiling of three psychiatric disorders. Genome Med. 9, 72 (2017).

83. Johnson, E. K., Matkovich, S. J. & Nerbonne, J. M. Regional Differences in mRNA and lncRNA Expression Profiles in Non-Failing Human Atria and Ventricles. Sci. Rep. 8, 13919 (2018).

84. Gupta, R. et al. Comparing in vitro human liver models to in vivo human liver using RNA-Seq. Arch. Toxicol. 95, 573–589 (2021).

85. Boegel, S. et al. HLA typing from RNA-Seq sequence reads. Genome Med. 4, 102 (2012).

86. Reynisson, B., Alvarez, B., Paul, S., Peters, B. & Nielsen, M. NetMHCpan-4.1 and NetMHCIIpan-4.0: improved predictions of MHC antigen presentation by concurrent motif deconvolution and integration of MS MHC eluted ligand data. Nucleic Acids Res. 48, W449–W454 (2020).

87. Yao, T. et al. Long-Read Sequencing Reveals Alternative Splicing-Driven, Shared Immunogenic Neoepitopes Regardless of SF3B1 Status in Uveal Melanoma. Cancer Immunol. Res. 11, 1671–1687 (2023).

88. You, Y. et al. Benchmarking long-read RNA-sequencing technologies with LongBench: a cross-platform reference dataset profiling cancer cell lines with bulk and single-cell approaches. Preprint at 10.1101/2025.09.11.675724 (2025).

89. Liu, Q. et al. LongGF: computational algorithm and software tool for fast and accurate detection of gene fusions by long-read transcriptome sequencing. BMC Genomics 21, 793 (2020).

90. Li, H. New strategies to improve minimap2 alignment accuracy. Bioinformatics 37, 4572–4574 (2021).

91. Frankish, A. et al. GENCODE 2021. Nucleic Acids Res. 49, D916–D923 (2021).

92. Cotto, K. C. et al. Integrated analysis of genomic and transcriptomic data for the discovery of splice-associated variants in cancer. Nat. Commun. 14, 1589 (2023).

93. Aguet, F., et al. The GTEx Consortium atlas of genetic regulatory effects across human tissues. bioRxiv 787903 (2019) doi:10.1101/787903.

94. Li, G. et al. A pan-cancer atlas of T cell targets. 2025.01.22.634237 Preprint at 10.1101/2025.01.22.634237 (2025).

95. Chen, J. et al. Targeting CLDN18.2 in cancers of the gastrointestinal tract: New drugs and new indications. Front. Oncol. 13, (2023).

96. Carstens, E. J. et al. Modeling and addressing on-target/off-tumor toxicity of claudin 18.2 targeted immunotherapies. Nat. Commun. 16, 9651 (2025).

97. Tio, D. et al. Expression of cancer/testis antigens in cutaneous melanoma: A systematic review. Melanoma Res. 29, 349–357 (2019).

98. Hofmann, O. et al. Genome-wide analysis of cancer/testis gene expression. Proc. Natl. Acad. Sci. U. S. A. 105, 20422–20427 (2008).

99. Karczewski, K. J. et al. The mutational constraint spectrum quantified from variation in 141,456 humans. Nature 581, 434–443 (2020).

100. Wickland, D. P. et al. Comprehensive profiling of cancer neoantigens from aberrant RNA splicing. J. Immunother. Cancer 12, e008988 (2024).

101. Gabrusiewicz, K. et al. Glioblastoma-infiltrated innate immune cells resemble M0 macrophage phenotype. JCI Insight 1, e85841, 85841 (2016).

102. Bao, Z.-S. et al. RNA-seq of 272 gliomas revealed a novel, recurrent PTPRZ1-MET fusion transcript in secondary glioblastomas. Genome Res. 24, 1765–1773 (2014).

103. Lavallée, V.-P. et al. RNA-sequencing analysis of core binding factor AML identifies recurrent ZBTB7A mutations and defines RUNX1-CBFA2T3 fusion signature. Blood 127, 2498–2501 (2016).

104. Goldmann, T. et al. PD-L1 amplification is associated with an immune cell rich phenotype in squamous cell cancer of the lung. Cancer Immunol. Immunother. CII 70, 2577–2587 (2021).

105. Heeke, S. et al. Tumor- and circulating-free DNA methylation identifies clinically relevant small cell lung cancer subtypes. Cancer Cell 42, 225–237.e5 (2024).

106. Sun, Q. et al. Long-read sequencing reveals the landscape of aberrant alternative splicing and novel therapeutic target in colorectal cancer. Genome Med. 15, 76 (2023).

107. Lesluyes, T. et al. Genomic and transcriptomic comparison of post-radiation versus sporadic sarcomas. Mod. Pathol. Off. J. U. S. Can. Acad. Pathol. Inc 32, 1786–1794 (2019).

108. Glinos, D. A. et al. Transcriptome variation in human tissues revealed by long-read sequencing. Nature 608, 353–359 (2022).

109. Reese, F. et al. The ENCODE4 long-read RNA-seq collection reveals distinct classes of transcript structure diversity. bioRxiv 10.1101/2023.05.15.540865 (2023) doi:10.1101/2023.05.15.540865.

110. Xie, N. et al. Neoantigens: promising targets for cancer therapy. Signal Transduct. Target. Ther. 8, 1–38 (2023).

111. Apavaloaei, A. et al. Tumor antigens preferentially derive from unmutated genomic sequences in melanoma and non-small cell lung cancer. Nat. Cancer 6, 1419–1437 (2025).

112. Kwok, D. W. et al. Tumor-wide RNA splicing aberrations generate immunogenic public neoantigens. bioRxiv 10.1101/2023.10.19.563178 (2023) doi:10.1101/2023.10.19.563178.

113. Karasikov, M. et al. Efficient and accurate search in petabase-scale sequence repositories. Nature 1–9 (2025) doi:10.1038/s41586-025-09603-w.

114. Oka, M. et al. Aberrant splicing isoforms detected by full-length transcriptome sequencing as transcripts of potential neoantigens in non-small cell lung cancer. Genome Biol. 22, 9 (2021).

115. Martin, M. V. et al. The Neo-Open Reading Frame Peptides That Comprise the Tumor Framome Are a Rich Source of Neoantigens for Cancer Immunotherapy. Cancer Immunol. Res. 12, 759–778 (2024).

116. Ji, S. et al. Large-scale transcript variants dictate neoepitopes for cancer immunotherapy. Sci. Adv. 11, eado5600 (2025).

